# Starve-Feed Cycles Direct Quiescence to Proliferation Transitions in *Drosophila* Follicle Stem Cells via Transcriptional Regulation

**DOI:** 10.64898/2026.01.22.701111

**Authors:** Eric H. Lee, Jacqueline C. Simonet, Daniel Zinshteyn, Zhen Fu, Saranya Ananth, Catharine Wingle, Ebony R. Dyson, Brendan D. Russell, Ruthie M. Njagi, Gabrielle N. Stills, Damiya Ringgold, Aminah Johnson, Arslie Louis-Jacques, Gabriel Vaughn, Maxwell Saurman, Dara M. Ruiz-Whalen, Jennifer I. Alexander, Yan Zhou, Alana M. O’Reilly

## Abstract

Stem cell quiescence is a reversible state in which cells temporarily exit the cell cycle but remain poised to reenter, on cue. Robust protection of the balance between stem cell quiescence and proliferation (Q->P) is critical for long-term tissue health. Proliferation without proper resources drives disease states including cancer or birth defects. Conversely, extended periods of quiescence can lead to irreversible senescence, causing stem cell loss, aging symptoms, and vulnerability to oncogenic transformation. Diet is a central regulator of Q->P. Tissue stem cells are particularly impacted, entering periods of quiescence during nutrient restriction, with rapid induction of proliferation upon feeding. We demonstrated previously that the Hedgehog (Hh) signaling pathway is necessary and sufficient for controlling Q->P responses to dietary changes in epithelial Follicle Stem Cells (FSCs) in the fly ovary. The Hh effector, Cubitus Interruptus (Ci), is a transcriptional regulator that mediates the feeding response. To identify Ci-induced Q->P regulators, we labeled transcripts that are induced in FSCs during the 6-hour Q->P timecourse using thiouracil tagging (TU-tagging), and sequenced TU-tagged messages versus Input to prioritize candidates. Unexpectedly, cell cycle regulators were not induced, suggesting that other mechanisms control Q->P in FSCs. We describe a sequential screening approach that uncovered seven novel, feeding-dependent Q->P regulators, including a cholesterol transporter and, surprisingly, glial and neuronal regulators. Our results highlight the importance of dynamic regulation of gene expression for translation of dietary signals by stem cells, uncovering new pathways for mechanistic investigation.

## Introduction

Diet is a fundamental, but inherently tumultuous, regulator of stem cells^1^. Periods of feeding and fasting promote dramatic changes in stem cell state, with low nutrient conditions ensuring quiescence, and feeding rapidly inducing proliferation to enable tissue generation or repair. The importance of feed-fast cycles at the organismal level has begun to emerge, with studies in humans and animals demonstrating health benefits of continuous or intermittent dietary restriction to reduce metabolic disease and, remarkably, to extend lifespan^2, 3^. Our goal is to define the molecular mechanisms that mediate feeding-dependent quiescence to proliferation (Q->P) transitions in stem cells, as a critical step towards development of interventions that can benefit organismal health.

Housed in microenvironments called niches, stem cells rely on their surroundings to control the fate decision between maintaining stem cell identity or differentiating^4^. In cases like *Drosophila* germline stem cells (GSCs), signals from the niche ensure long functional lifespan of individual GSC clones and inheritance of stem cell function through generations^5^. Other stem cells exist in an aggressive, competitive environment, where limited niche space drives stem cell selection, and losers of the competition are displaced to undergo differentiation ^6-9^. Although high proliferative rates are associated with retention^10-19^, hypercompetition drives clonality in stem cell pools over time^20^, a characteristic of stem cell loss and aging. Thus, strategies that maintain balance via Q->P transitions may prove effective for long-term health of the stem cell pool^21, 22^

In our previous work, we demonstrated that epithelial Follicle Stem Cells (FSCs) in the *Drosophila* ovary enter quiescence upon nutrient restriction and re-enter the cell cycle rapidly upon feeding^23^. This process depends on ingestion of cholesterol, which triggers a multi-step mechanism that leads to release of the Hedgehog (Hh) ligand from the Hh-producing terminal filament and cap cells at the anterior tip of the ovary (Fig. 1A)^23^. Hh induces the Q->P transition by inhibiting its receptor, Patched (Ptc), on the surface of FSCs, releasing the downstream effector, Smoothened (Smo), to activate Cubitus Interruptus (Ci), a homolog of the mammalian Gli family of transcription regulators^23^. Dividing FSCs predominantly adopt one of two fates in the short term. Daughter cells that remain in the niche maintain their stem cell identity. Conversely, daughter cells that exit the niche to the posterior initiate differentiation, adopting a follicle cell fate^13, 24-26^. Follicle cells coalesce into a single layered cuboidal epithelium that encapsulates developing germ cell cysts, creating follicles called “egg chambers” that develop through 14 iterative stages to produce mature eggs^27-29^.

**Figure 1:**
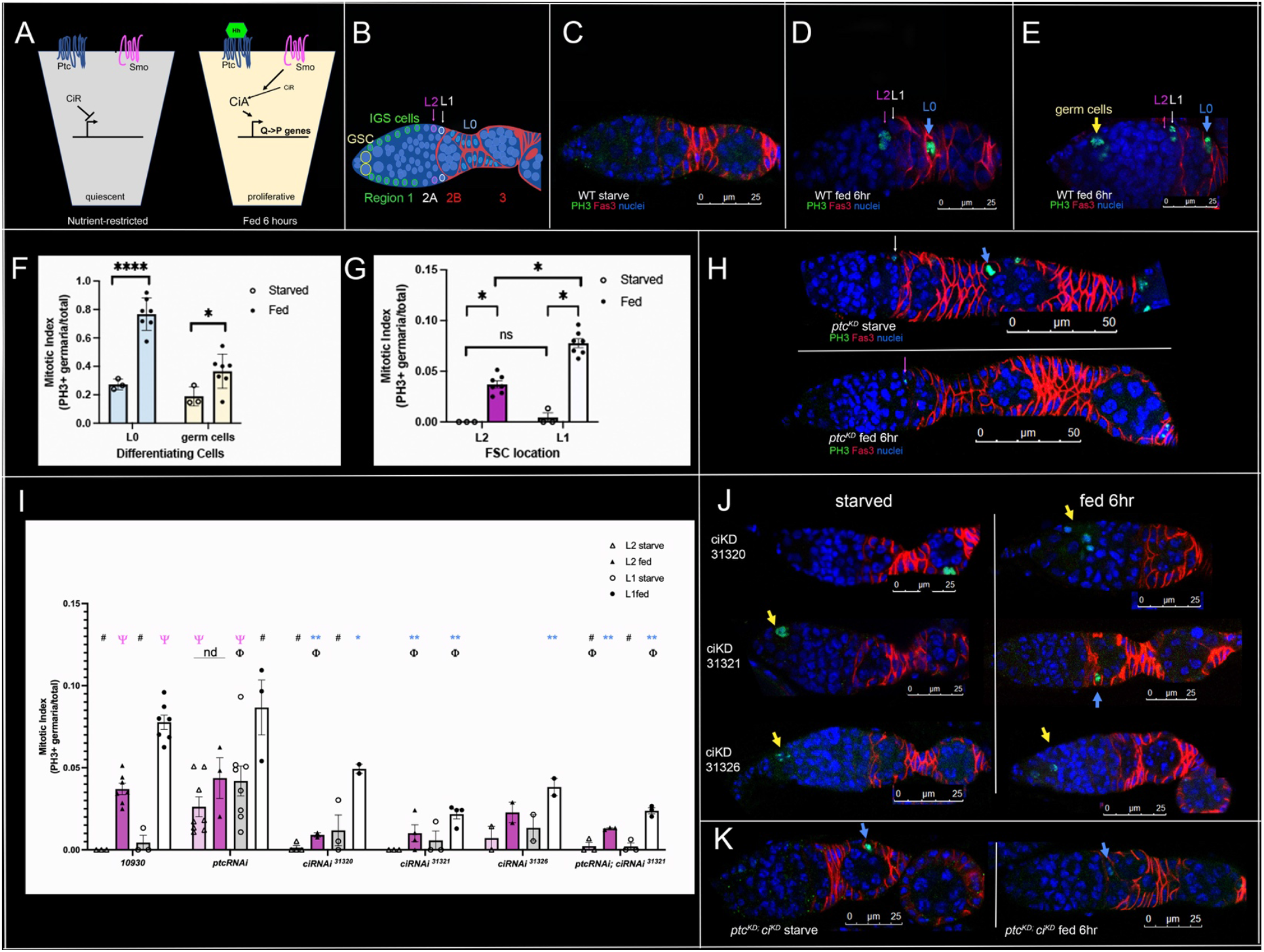
Hh-mediated transcription is required for Q->P transitions in FSCs. A) Model for feeding-dependent activation of Hh signaling. In quiescent FSCs, Hh ligand is absent, Ptc suppresses Smo, and the repressor version of Ci (CiR) is actively repressing Hh-dependent transcripts. Upon feeding, cholesterol induces Hh release, relieving Ptc-mediated Smo inhibition and activating downstream transcriptional activation by the activator form of Ci (CiA). B) Schematic of the germarium, with Regions 1-3 indicted (bottom). IGS cells (green), follicle cells (red) and germline stem cells (GSCs, yellow) are indicated. Layer 1 (L1 white) and Layer 2 (pink) FSCs reside anterior to the Fas3 border. C) Nutrient-restricted (starved) control germaria exhibit no FSCs in mitosis. D) 6 hours after feeding, L2 FSCs (pink arrow) are labeled with anti-phosphohistone H3 (PH3, green), indicating mitosis. E) L1 FSCs (white arrow) labeled with PH3 (green) 6 hours after feeding. F, G) Quantitation of the mitotic index (PH3+ germaria/total) for (F) Layer 0 Fas3+ follicle cells (L0, light blue) and germ cells (yellow) and (G) Layer 2 (L2, pink) and Layer 1 (L1, white) FSCs. Significant increases in mitotic index from nutrient-restricted (starved) to fed are observed for all cell types (**** p<0.0001, *p<0.05). H) Nutrient-restricted (starved) versus fed *ptc*^*KD*^ germaria indicating mitotic L1 (starved, white arrow) or L2 (fed, pink arrow) FSCs. I) Quantitation of the mitotic index (PH3+ germaria/total) for Layer 2 (starved: light pink; fed: dark pink) and Layer 1 (starved: gray; fed: white) FSCs. Genotypes are indicated. 4′ indicates significant differences (p<0.05) vs. 10930 Layer-specific starved controls. * (p<0.05) and ** (p<0.01) indicate significant differences between fed *10930* controls and Layer-specific fed candidate genes. **#** indicates significant differences relative to Layer-specific *ptc*^*KD*^ starved conditions. Φ indicates significant differences (p<0.05) versus Layer-specific, fed *ptc*^*KD*^ controls. nd highlights lack of significance between *ptc*^*KD*^ starved and fed conditions in Layer 2.

The essential role of Ci in mediating the feeding-induced Q->P transition^23^ implicates dynamic changes in gene expression as a critical element of FSC homeostasis. Recently, multiple single cell RNA sequencing studies have uncovered cell-specific ovary transcripts^30-35^. Matching expression patterns with known markers has been leveraged to create candidate gene signatures for particular subsets of cells in the stem cell compartment of the ovary, called the germarium (Fig. 1B). Clear delineation of germ cells from somatic cells suggests there may be opportunities for refinement of cellular identity using transcriptional profiling^30, 31, 33^. For example, several studies have mapped distinctions between geographically distinct populations of Inner Germarial Sheath Cells (IGS cells, also called Escort Cells) that support early germline development (Fig. 1B)^36^, finding changes in gene signature in IGS cells located in anterior, central, or posterior locations in the germarium^31, 33-35^. Likewise, gene signatures for specific stages of germ cell development or distinctions between somatic polar versus stalk cells can be clearly described^30, 33, 34^. Granular distinctions, for example between very early germ cells (Germline Stem Cells (GSCs) versus cystoblasts), have been more challenging given the extensive overlap in gene expression between these cell types, despite clear evidence of different functions^27, 30, 33, 36^. Along these lines, there remains substantial disagreement regarding the gene signature of FSCs, which express markers that overlap to varying degrees with IGS cells to the anterior and follicle cells to the posterior. Whereas some groups have proposed anchor markers for FSCs^31, 35^, none have identified a clearly unique FSC marker or signature^30, 33, 34^.

Despite these extensive efforts to pin down FSC identity with molecular and spatial precision, a more exciting possibility is that FSCs adapt to instructional signals that reflect the needs of the tissue in real time. Consistent with this idea, we recently demonstrated that key markers of FSCs and follicle cells change dramatically during the transition from quiescence to proliferation^37^. Another study highlighted changing functional capacity of FSCs to produce clonally derived follicle cell daughters in response to starving and feeding^31^, consistent with our findings. Moreover, proliferation rates and differentiation character change depending on FSC position and signaling gradients^38, 39^, with rapid switching between locations or genetic alteration of signaling gradients influencing FSC behavior and proliferation in real time^13, 26, 38, 39^. Thus FSCs may confound delineation of permanent, defining gene signatures, instead highlighting the potential of this system to uncover combinations of factors that are induced or suppressed in response to rapidly changing tissue demands to control stem cell behaviors.

Here, we used sequential screening to uncover feeding-dependent transcriptional regulators of Q->P transitions in FSCs. Our first step was to endogenously label nascent transcripts induced within FSCs during the 6 hour Q->P transition timecourse using Thiouracil Tagging (TU-Tagging)^40, 41^. We then purified and sequenced transcripts that were differentially enriched in FSCs and early follicle cells whose expression changed during the 6 hour Q->P transition. Next, we evaluated expression of prioritized candidate genes in the germarium using the RNA *in situ* hybridization approach, RNAScope^42^, pinpointing genes enriched in FSCs, specifically. Finally, we performed functional analysis of genes isolated via our primary and secondary gene expression screens to identify novel regulators of Q->P. We present evidence that feeding-induced, cell autonomous Q->P regulators are found in predictable pathways, including cholesterol trafficking and feeding regulation, and also in unexpected pathways that regulate neuronal and glial signaling. This work highlights the value of conducting unbiased, forward genetic screening for identification of dynamic new factors critical for stem cell regulation.

## RESULTS

The Hh pathway translates nutritional signals to promote Q->P transitions in FSCs (Fig. 1A)^23^. FSCs fail to proliferate in nutrient-restricted wild-type flies raised on grape juice plates, which lack any source of protein or fat^23^. FSCs remain in this state for as long as the flies are maintained on grape juice plates^23^. Proliferation is induced in FSCs upon feeding, with yeast providing a robust source of nutrients necessary for promoting Q->P transitions. FSCs reside in a multicellular structure called the germarium, found at the anterior end of the ovary (Fig. 1B). The germarium houses both FSCs and germline stem cells (GSCs), which produce 16-cell germline cysts that undergo 14 developmental stages to form a mature egg. Interactions of germline cysts with somatic cells define developmental regions of the germarium, with Region 1 housing the germline stem cells and the somatic cells that comprise their niche. Region 2 is divided into Region 2A and Region 2B, based on interactions of germline cysts with stationary Inner Germarial Sheath (IGS) cells in Region 2A or follicle cells in Region 2B (Fig. 1B). FSCs reside in between these two somatic cell populations, at the border between Region 2A and 2B. Shifts in the relationship between germline cysts and somatic cells during development, coupled with major shape changes that occur as germline cysts traverse the germarium, make use of the Region 2A/2B border challenging as a definitive marker of FSC location. Luckily, the cell membrane protein Fasciclin 3 (Fas3) marks differentiating follicle cells, clearly delineating FSCs from their differentiating daughters. FSCs reside one and two cell diameters anterior to the Fas3 border, referred to as Layer 1 and Layer 2, respectively (Fig. 1B). Layer 1 and Layer 2 are coincident with the Region 2A/2B border (Fig. 1B), providing both developmental and molecular markers to consistently pinpoint the location of FSCs.

FSCs in Layers 1 and 2 can interchange, but their relative anterior-posterior position dictates key parameters including proliferation rate^37, 38, 43, 44^. Layer 2 cells proliferate less frequently than Layer 1 cells^38, 39^, and express lower levels of the differentiation markers Eyes Absent (Eya) and Castor (Cas) relative to Layer 1 cells^37^. Both proliferation and differentiation marker expression are affected by diet, with proliferation arrest coupled with reduction or loss of expression of Eya and Cas observed under nutrient-restriction conditions; expression and cell division are restored within 6 hours after feeding (Fig. 1C^23, 37^). Feeding stimulates cell cycling, with mitotic (phosphohistone H3 positive (PH3+)) FSCs observed in Layer 2 (Fig. 1D) or Layer 1 (Fig. 1E) by the 6 hour timepoint. Thus, FSCs present an ideal model of epithelial stem cells for delineation of the mechanisms that control Q->P transitions in response to feeding.

To establish quantitative, baseline impacts of feeding on cells in different locations within the germarium, we scored the mitotic index, defined as germaria with at least one PH3+ cell, for 1) cells expressing Fas3 on all borders (Layer 0), 2) Layer 1 FSCs located adjacent to Fas3 positive cells to the anterior, 3) Layer 2 FSCs, located one cell anterior to Layer 1, and 4) germ cells (Fig. 1B-E). The mitotic index of Layer 0 (Fas3+) cells was about one third in nutrient-restricted conditions versus 6 hours after feeding (Fig. 1F, Table S1). A lower average mitotic index was observed in nutrient-restricted germ cells relative to post-feeding, with more variability in both the nutrient-restricted and fed pools (Fig. 1F, Table S1). These results demonstrate that differentiating cells can divide even in the absence of key nutrients, albeit at a substantially lower rate. By contrast, no mitotic cells in Layer 1 or Layer 2 were observed in nutrient-restricted conditions (Fig. 1G, Table S1), supporting the notion that FSCs in both Layers enter a state of quiescent arrest in the absence of key nutrients. Mitotic rates differed between Layer 1 and Layer 2 cells 6 hours after feeding, with Layer 1 cells proliferating approximately twice as frequently (Fig. 1G, Table S1), consistent with reported differences in proliferation rates of 1.35 to 3-fold between Layers 2 and 1^38, 39^.

We next compared the impacts of altering Hh signaling on mitotic index in the same four locations. Hh activates downstream effectors by inhibiting the action of its receptor, Patched (Ptc) (Fig. 1A). Hh-bound Ptc is unable to suppress the activity of Smoothened (Smo), enabling signaling to proceed (Fig. 1A)^45^. Smo functions to promote accumulation of the transcriptional activator Ci (CiA), which induces Hh-dependent gene targets to elicit cellular responses (Fig. 1A). Thus, loss of *ptc* expression constitutively activates Hh signaling^45, 46^. We expressed RNAi targeting *ptc* using the *109-30*-Gal4 driver, which drives expression in FSCs and early follicle cells^23, 47^. As expected, *ptc* knockdown (*ptc*^*KD*^) in FSCs and early follicle cells had no effect on the mitotic index in germ cells, which do not express *ptc*^*RNAi*^, serving as an internal control (Fig. S1). Mitosis of *ptc*^*KD*^ cells in Layers 0,1, or 2 was the same as wild-type 6 hours after feeding (Fig. 1H, I, Fig. S1, Table S1), indicating that loss of *ptc* had no additional effect on FSC mitotic index in response to feeding. This result is consistent with the model that feeding-induced signals rely on Hh-mediated inhibition of Ptc. By contrast, *ptc*^*KD*^ dramatically affected the mitotic index of cells in Layers 1 and 2, as well as differentiating follicle cells in Layer 0, under nutrient-restriction conditions (Fig. 1H,I, Fig S1, Table S1)). Instead of entering quiescence like nutrient-restricted wild-type FSCs, which were never observed in mitosis, *ptc*^*KD*^ FSCs divided, with a mitotic index that was 66% (Layer 2), 69% (Layer 1), and 79% (Layer 0) of that observed for fed wild-type FSCs (Fig. 1I, Table S1). These results support the notion that inactivation of Ptc upon feeding is responsible for Q->P transitions in FSCs, and demonstrate that reducing *ptc* expression bypasses the feeding requirement for Q>P induction.

Our previous results demonstrated that the transcriptional Hh effector, *ci*, is necessary for Q->P transitions in FSCs^23, 45, 47^, suggesting that induction or suppression of expression of Q->P regulatory genes is the critical mechanism for triggering Q->P in response to feeding. If this hypothesis is correct, we expected that *ci* knockdown (*ci*^*KD*^) would 1) block feeding-induced Q->P transitions and 2) suppress proliferation induction by *ptc*^*KD*^ under nutrient restriction or fed conditions. Both of these predictions are supported by our data for Layer 2 FSCs, with a reduction in mitotic index in *ci*^*KD*^ cells of >3-fold relative to fed wild-type controls (Fig. 1I,J, Table S1). Impacts on fed Layer 1 FSCs were less severe, ranging from 1.5-2.25 fold reduced mitotic index in *ci*^*KD*^ cells relative to fed wild-type FSCs (Fig. 1I,J, Table S1). No significant change was observed for Layer 0 cells (Fig. S1), suggesting that Hh functions preferentially in FSCs. Reducing *ci* expression fully suppressed proliferation in nutrient-restricted or fed *ptc*^*KD*^ Layer 2 and Layer 1 FSCs (Fig. 1I, K, Table S1), demonstrating that transcription is required for FSC proliferation under conditions where Hh signaling is constitutively active, regardless of feeding status.

### The feeding-dependent transcriptome

Given the critical role of *ci*-dependent transcription in Q->P regulation, we sought to define the FSC-specific changes in gene expression that occur over a 6 hour timecourse after feeding. Transcripts newly synthesized during Q->P transitions were labeled using the TU-tagging technique (Fig. 2A^40, 41^). To do this, the yeast enzyme UPRT was expressed under control of *109-30-*Gal4 *tub*Gal80^ts^, which drives expression of genes containing UAS promoter elements in FSCs and early follicle cells in a temperature-controlled manner. At the restrictive temperature (18°C), flies were fed 4-thiouracil (TU), the substrate for UPRT. After switching to the permissive temperature to promote UPRT expression, flies were subjected to a brief period (3 days) of protein and lipid restriction in the presence of TU, driving FSCs into quiescence. Feeding with yeast paste + TU then induced Q->P^56 68^. Our previous work demonstrated that peak mitosis occurs in FSCs 6 hours after feeding^23, 37^. We reasoned that genes required for cell cycle entry would be induced prior to the mitotic peak and maintained at the 6 hour timepoint. TU-containing transcripts from nutrient-restricted or fed flies, 3 or 6 hours after feeding, were biotinylated, streptavidin-precipitated and subjected to RNA sequencing (referred to as “TU-tagged” going forward). Input RNA from the same samples was also sequenced, serving as a control for transcripts that changed generally, but were not specific to FSCs or early follicle cells. Using standard criteria of a 2-fold change and false discovery rate (FDR) of 5%, feeding-dependent genes were clustered into three groups: 1) “Background” genes whose expression changed in the pool of Input RNA from nutrient-restricted to fed conditions (green, Fig. 2B), 2) “Enriched” genes with higher expression in the streptavidin-precipitated pool versus the Input pool at the fed timepoint (blue, Fig. 2B), and 3) “Target” genes that changed expression from nutrient-restriction to fed within the TU-tagged pool (red, Fig. 2B). Cumulatively, 2161 genes changed expression between nutrient-restriction and the three hour timepoint, with 1425 genes changing at the 6 hour timepoint, highlighting the dramatic changes in gene expression that occur in both TU-tagged and Input transcripts in response to feeding.

**Figure 2:**
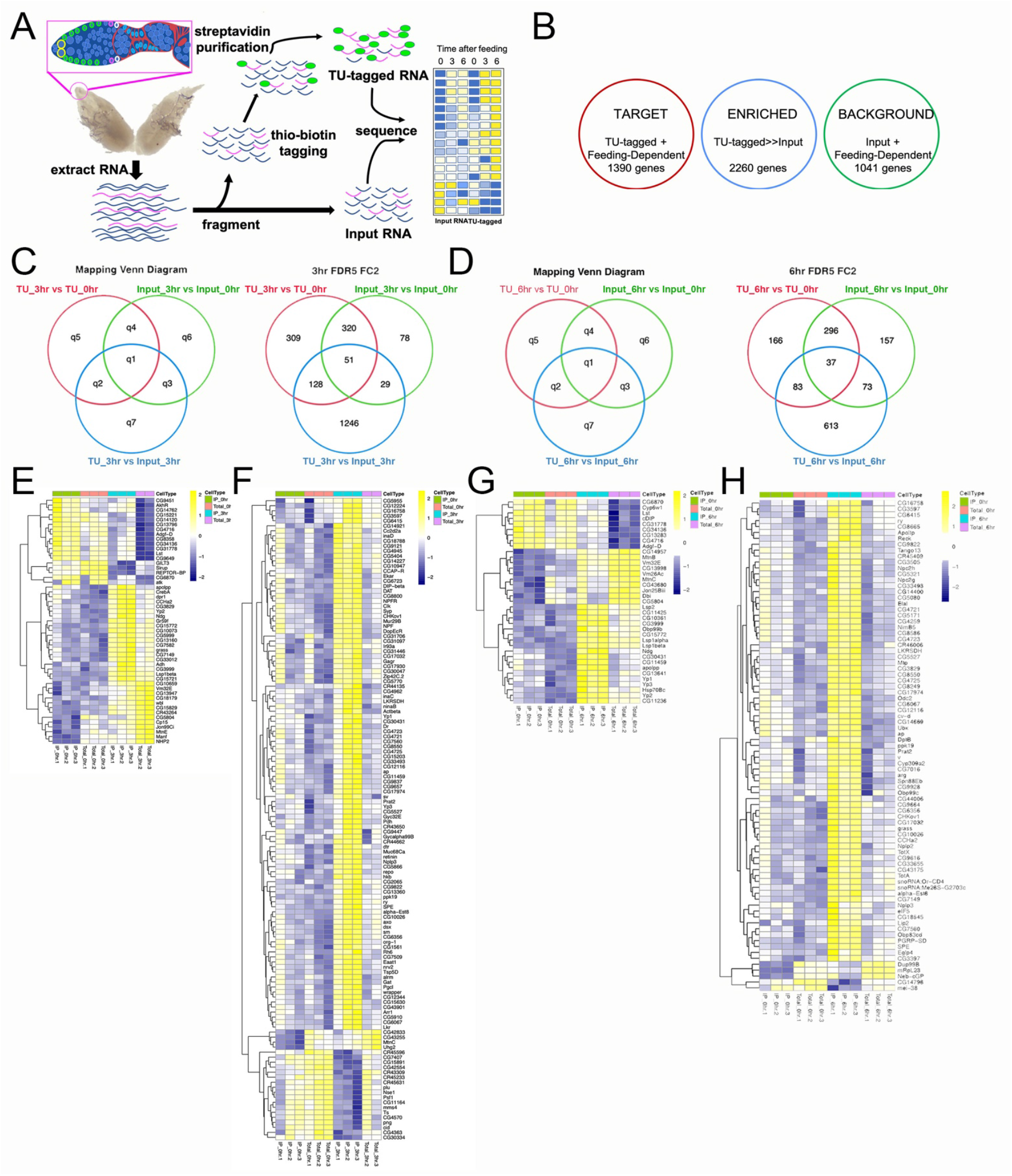
TU-tagging uncovers 300 known genes as Q->P regulatory candidates. A) Schematic of TU-tagging approach. Thiouracil-labeled RNA is precipitated with biotin-streptavidin. TU-tagged and Input RNA are sequenced, generating heat maps of gene expression before (time 0) and after feeding (3, 6 hours). B) Three gene clusters were used for the analysis: a “Target” pool of TU-tagged genes that changed gene expression 3 or 6 hours after feeding (red), an “Enriched” pool in which TU-tagged messages were at expressed at least 2-fold higher than Input messages (blue), and a “Background” pool of genes that changed expression over time in the Input, reflecting changes that are not specific to FSCs. C,D) Venn diagrams indicate numbers of genes in each of 7 clusters (q1-q7) 3 hours after feeding (C) or 6 hours after feeding (D). E-H) Heat maps of genes in q1 (E, G) or q2 (F, H) 3 hours (E, F) or 6 hours (G, H) after feeding.

We focused on defining genes that mediate the Q->P transition in FSCs. Extensive prior work in many systems identified transcriptional induction of the cell cycle regulators Cyclin E and Cdk2 as critical triggers for Q->P transitions^48-50^. Notably, Cyclin E is a known Ci target gene^51-53^ that can influence FSC proliferation^38, 39^. We evaluated enrichment over time (TU-tagged/Input at 0, 3, and 6 hours after feeding) of 20 cell cycle regulators (Fig. S2). No significant changes were observed during the 6-hour timecourse (Fig. S2). As such, other transcription-dependent mechanisms must promote Q->P in FSCs.

To uncover genes enriched in streptavidin-precipitated pools that change over time, Venn diagrams were created using Target (red, TU-tagged, feeding-dependent), Enriched (blue, higher expression in TU-tagged versus Input), and Background (green, feeding-dependent changes within the Input) gene clusters. Clusters were labeled q1-q7, with single parameters (q5, q6, q7) given lower priority and overlapping parameters (q1, q2, q3, q4) considered of higher interest (Fig. 2C,D). We chose to focus on candidate Q->P regulators in Clusters q1 and q2, which 1) changed expression in response to feeding and 2) were enriched in streptavidin-precipitated, TU-tagged pools relative to Input mRNA. Genes in q1 increased in both TU-tagged and Input pools, but were enriched in the TU-tagged pool. Genes in q2 increased in response to feeding only in the TU-tagged pool. 179 FSC-enriched genes were induced 3 hours after feeding (Aq1 and Aq2, Fig. 2C), and 120 were induced 6 hours after feeding (bq1 and bq2, Fig. 2D) (fold change >2, FDR<0.05). This Q->P gene list included 142 previously characterized and named genes (85 at 3 hours, 57 at 6 hours) as well as 157 genes that have not yet been named or studied in detail, referred to by their so-called “CG” numbers (Fig. 2E-H).

### FSC-enriched expression

Our primary focus for this screening effort was to identify genes that mediate the Q->P transition. Since no cell cycle regulators were identified as feeding-dependent or enriched in *10930*-Gal4-expressing cells (Fig. 2B), we conducted an RNA *in situ* hybridization analysis to narrow our focus within the q1 and q2 clusters. To do this, we collected fly lines bearing insertion of the *Gal4* transcriptional activator in candidate gene loci and conducted RNAi *in situ* hybridization analysis using the RNAScope method^42^. Probes targeting mRNA for Fas3, the marker of differentiating follicle cells (Fig.1), and Gal4 were used for the analysis, with the idea that Gal4 patterning would reflect normal expression of the candidate gene into which it was inserted. To validate and optimize the approach, we conducted RNAScope using flies bearing Gal4 insertions in genes with known patterns in FSCs and germaria (Fig. 3). In flies lacking Gal4 altogether (*w*^*1118*^, Fig. 3A), no Gal4 signal was observed. By contrast, the probe highlighted *armadillo*-Gal4 expression in all cells of the germarium (Fig. 3B), consistent with its known functional roles in germline and somatic cells during early oogenesis^54-59^. *pCOG*-Gal4 enables Gal4 expression from the *otu* locus, with expression concentrated in germ cells (Fig. 3C)^60^. *castor*-Gal4 (*cas*-Gal4) exhibited strong expression in polar and stalk cells and weaker expression in FSCs and early follicle cells, consistent with its concentration-dependent role in promoting polar and stalk cell differentiation (Fig. 3D)^44, 61^. As expected, *109-30*-Gal4 flies, which bear Gal4 inserted in the *sickie* (*sick*) gene^37^, exhibited robust Gal4 expression in FSCs, with lower levels in early follicle cells and stalk cells (Fig. 3E)^23, 47, 62, 63^. Finally, *Wnt4*-Gal4, which has been proposed as a marker to define the FSC lineage^31^, was expressed relatively non-specifically, with higher levels in terminal filament and IGS cells, in addition to sporadic expression in FSCs (Fig. 3F). These results confirm previously reported expression patterns, correlating transcription with reporter activity or protein expression and supporting use of RNAScope as a secondary screening method.

**Figure 3:**
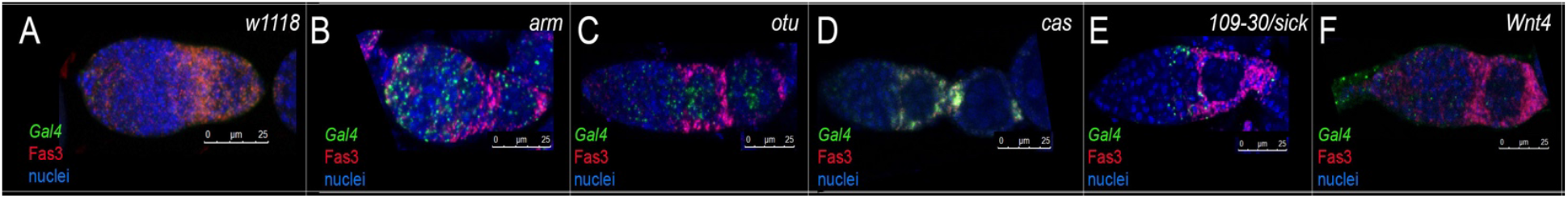
RNAScope reflects known patterns of genes expressed in the germarium. A-F) Germaria isolated from transgenic fly lines with Gal4 inserted into genes known to be expressed and functional in the germarium. Probes complementary to Gal4 (green) and Fas3 (red) were used for RNA in situ hybridization using the RNAScope approach. Nuclei are labeled (Draq5, blue). Scale bars are indicated for each image.

### Screen for FSC-enriched Gal4 expression

With RNAScope validated for analysis of gene expression in the germarium, we screened 46 fly lines with Gal4 inserted in previously characterized genes for expression in FSCs under steady-state feeding conditions, in which flies have unrestricted access to food. A group of 27 Gal4 lines exhibited either no expression or expression in many cell types, without obvious enrichment in FSCs (Fig. S3). Five additional lines exhibited concentrated expression in IGS cells, germ cells, follicle cells, terminal filament, or muscle cells, with no obvious enrichment in the FSC region (Fig. 4A). These 32 genes were eliminated from further analysis. 14 Gal4 lines exhibited expression that was enriched in the FSC niche region (Fig. 4B), albeit with additional expression in other cells in the germarium. Although FSC-exclusive expression was not observed, we reasoned that genes with enrichment in Layer 2 or Layer 1 cells might be important functionally for Q->P transitions.

**Figure 4.**
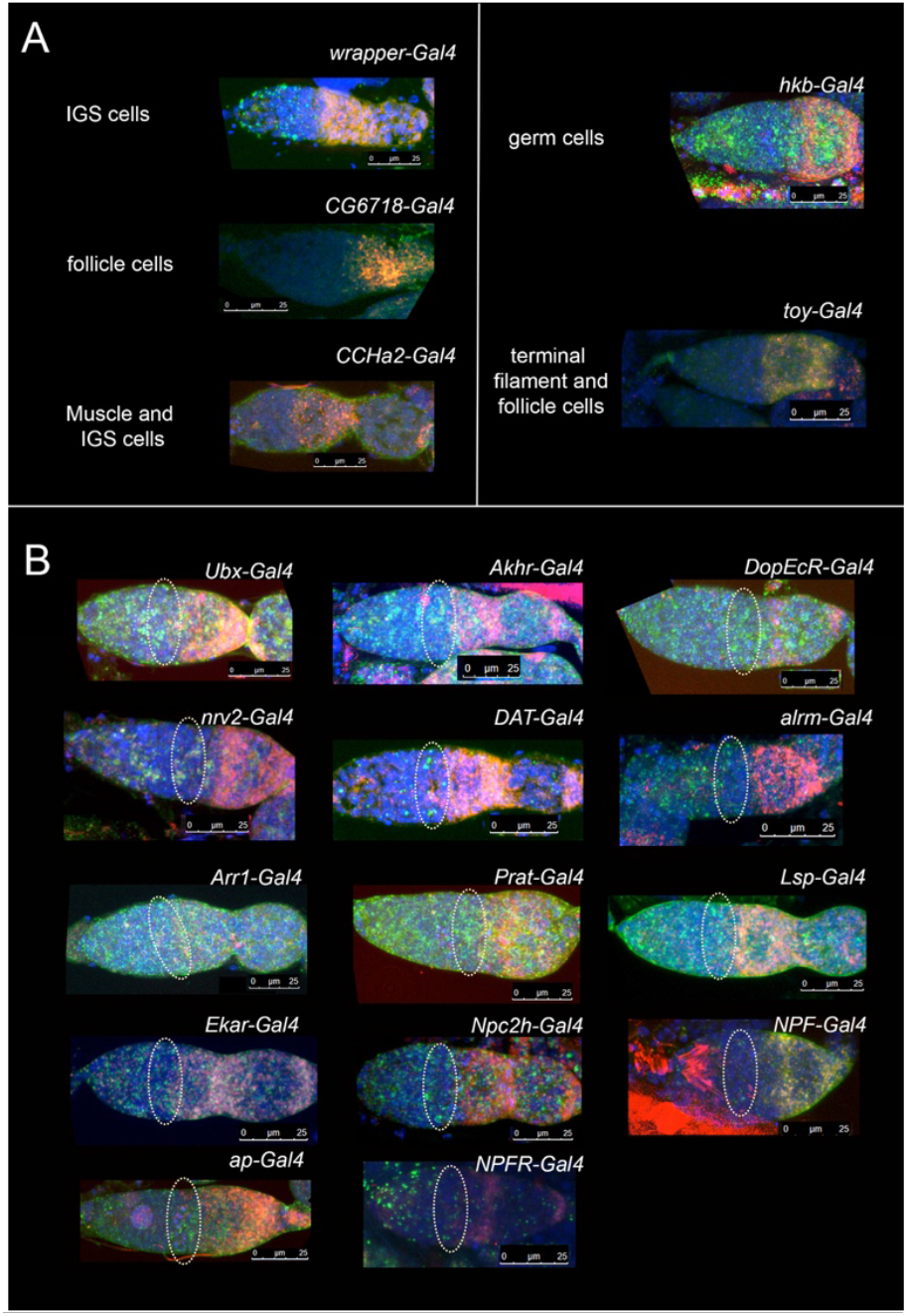
RNAScope of transgenic lines bearing insertion of Gal4 in Q->P regulatory candidates, using probes against Gal4 (green) and Fas3 (red). Nuclei are labeled (Draq5, blue). Candidates with Gal4 in cells other than FSCs (A), or enriched in FSCs (B) are shown. The FSC region is indicated by a dotted oval in B. Scale bars are indicated for each image.

### Feeding dependent induction of gene expression in FSCs

We next repeated our RNAScope analysis in germaria from nutrient-restricted versus 6-hour fed flies to determine which candidates exhibited robust induction of expression in FSCs upon feeding. Control Gal4 lines including *109-30*-Gal4, *arm*-Gal4, and *otu*-Gal4 exhibited similar levels and expression patterns under nutrient-restriction and 6-hour-fed conditions (Fig. 5A,F). *cas*-Gal4 exhibited diffuse, broad fluorescence in nutrient-restricted conditions, becoming punctate in the FSC region after feeding (Fig. 5A), consistent with changes in Cas expression at the protein level upon feeding^37^.

**Figure 5:**
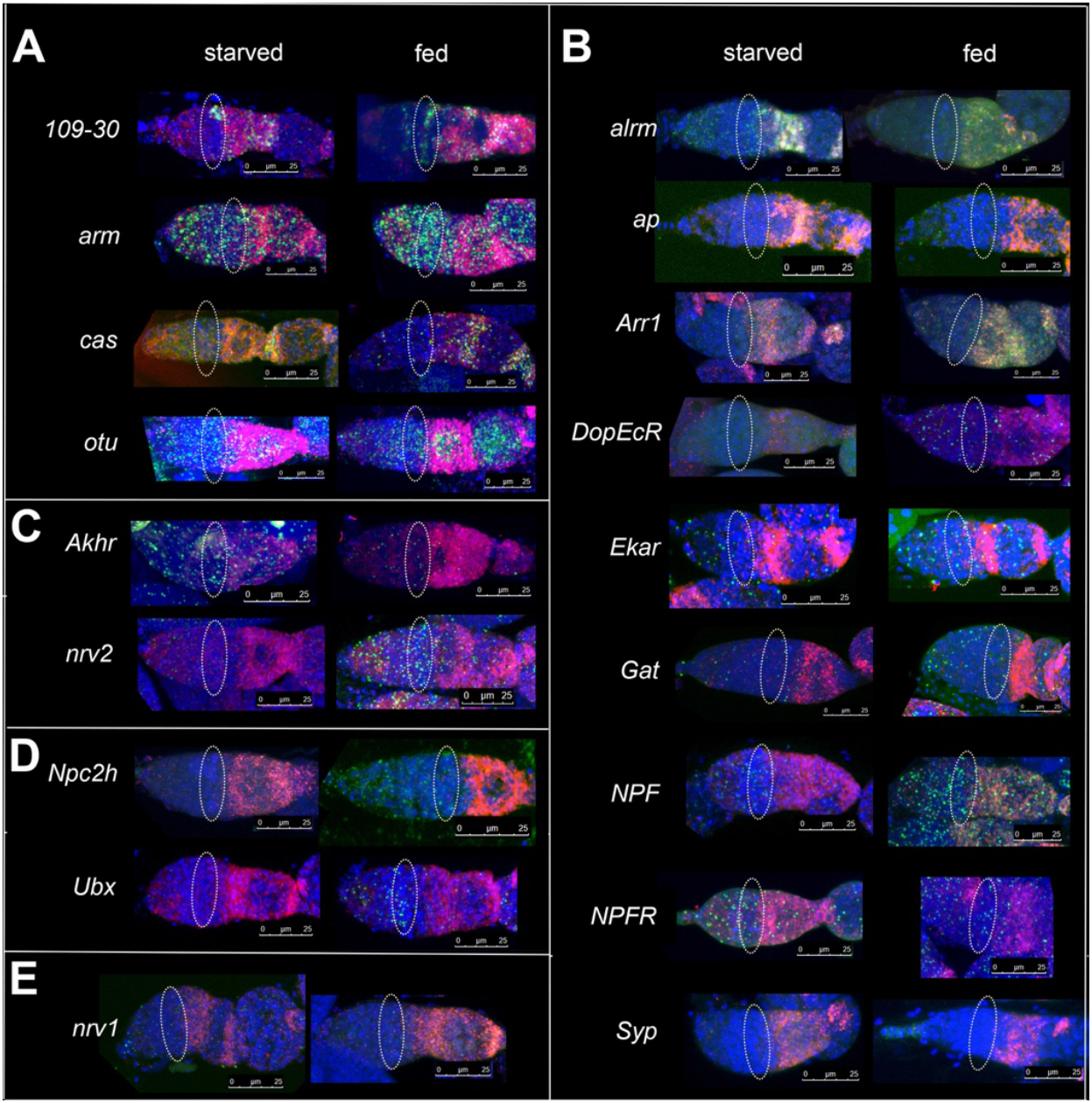
Feeding-dependence of gene expression. A-E) RNAScope of germaria bearing Gal4 expression in the indicated genes. Probes against Gal4 (green) and Fas3 (red) indicate gene expression in nutrient-restricted (“starved”) or 6-hours fed germaria. Nuclei are labeled (Draq5, blue). The FSC niche is outlined by a dashed oval. Scale bars are indicated for each image. A) Feeding-dependence of previously characterized genes. B) Candidates in the Aq2 cluster, which are enriched in the TU-tagged fraction and increase by 3 hours after feeding. C) Candidates in the Aq1 cluster, which are enriched in the TU-tagged fraction and increase in both the Input and the streptavidin-precipitated fraction by 3 hours after feeding. D) Candidates in the bq1 cluster, which are enriched in the TU-tagged fraction and increase in both the Input and the streptavidin-precipitated fraction by 6 hours after feeding. E) *nrv1-Gal4* germarium, a non-feeding-dependent example that shares function with *nrv2*, a feeding-dependent gene.

Next, we used RNAScope to narrow down candidates for functional analysis. We divided the candidates based on their RNAseq clusters, focusing on q1 and q2, as above. The largest group of candidates analyzed was in Aq2, genes that are induced 3 hours after feeding and enriched in TU-tagged, streptavidin-precipitated samples (Fig. 5B). The magnitude, pattern, and specificity of expression changes varied among the Aq2 gene pool (Fig. 5B). *Ekar*-Gal4 exhibited dramatically increased expression in the FSC region upon feeding, consistent with its identification as the strongest hit in a known gene by RNAseq (Fig. 2F). Additional candidates (*ap, Gat, NPF, NPFR*) indicated clear feeding-dependent FSC niche enrichment in a subset of germaria, supporting their candidacy for further investigation. In some cases, such as *Syp*-Gal4, the enhancer element captured by the Gal4 insertion may be specific for another cell type, as Gal4 signal was concentrated in terminal filament cells rather than FSCs (Fig. 5B). Genes classified as Aq1 (*Akhr, nrv2*, Fig. 5C), which were enriched in the TU-tagged fraction and induced in both Input and TU-tagged samples 3 hours after feeding, and bq2 (*Npc2h, Ubx*, Fig. 5D), which were induced 6 hours after feeding in the TU-tagged fraction, consistently exhibited substantial changes in expression from nutrient-restricted to fed conditions. Of the candidates tested, only *Akhr* exhibited reduced expression after feeding (Fig. 5C,F), consistent with the RNA sequencing results (Fig. 2C).

## Regulators of Q->P transitions

As a final step in our screen for Q->P regulators, we expressed RNAi targeting candidate genes under control of *10930*-Gal4 and measured the mitotic index in Layer 2 and Layer 1 FSCs and Layer 0 differentiating follicle cells relative to control wild-type FSCs. Our primary focus was on genes found to be 1) concentrated in FSCs in steady state feeding conditions, 2) absent or low expression in quiescent FSCs under nutrient-restriction, and 3) induced in FSCs upon feeding. Only one gene, the cholesterol transporter *Npc2h*, was necessary for feeding-stimulated proliferation in Layers 2, 1, and 0, indicating an essential role in mediating feeding signals to control proliferation induction in general (Fig. 6A-C, Table S1).

**Figure 6:**
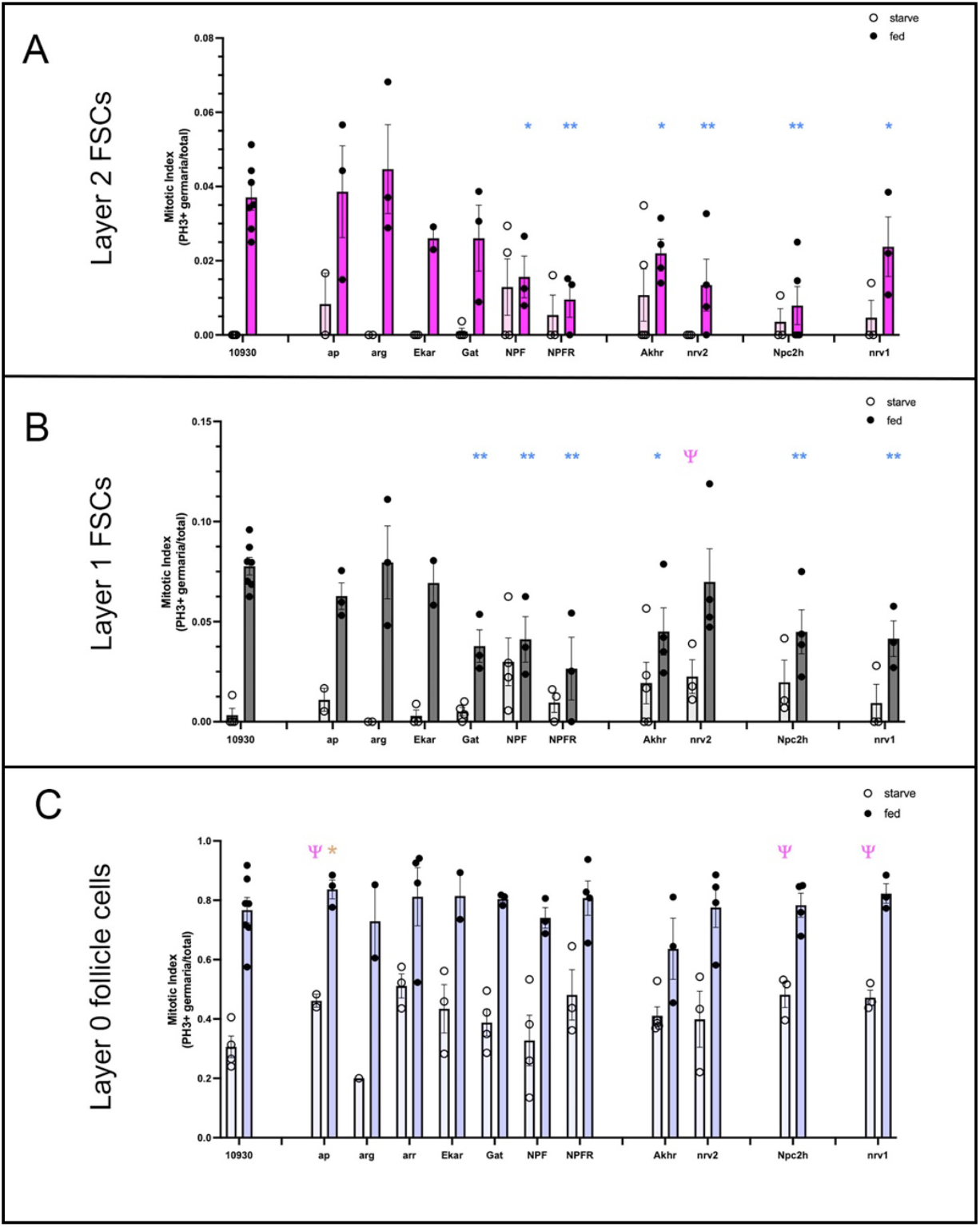
A subset of candidates are essential for reversible Q->P transitions. A-C) Quantitation of the mitotic index (PH3+ germaria/total) for Layer 2 (A, pink), Layer 1 (B, gray), or Layer 0 (C, blue) cells. Nutrient-restricted (“starved”, lighter colors, open circle) versus fed (darker colors, filled circle) conditions are shown for knockdown conditions for the genes indicated. * (p<0.05) and ** (p<0.01) indicate significant differences between fed *10930*Gal4 *tub*Gal80^ts^ (10930) controls and fed candidate genes. 4′ (p<0.05) indicates significant differences between starved 10930 controls and starved candidate genes. * indicates p=0.05. Values are detailed in the Supplementary Table.

In addition to cholesterol regulation, genes involved in neuronal and glial control scored as hits (Fig. 6, Table S1). Expression of the feeding hormone *NPF* was necessary for Q->P in Layer 2 and 1 FSCs, but had no effect on differentiating Layer 0 follicle cells (Fig. 6, Table S1). Consistent with the requirement for *NPF*, the *NPF Receptor* (*NPFR)* was necessary for Q->P in Layer 2 FSCs (Fig. 6, Table S1). *Akhr*, the receptor for the fly’s functional analog of glucagon, adipokinetic hormone (Akh), was also required for Q->P (Fig. 6, Table S1). Given that *Akhr* expression drops in response to feeding (Fig. 2C, 5C,F), we initially expected loss of *Akhr* to impact maintenance of quiescence. The unexpected requirement for feeding-dependent Q->P may indicate a metabolic transition phase that depends on Akhr-mediated metabolic processes utilized during quiescence, with other mechanisms driving the metabolic transition to steady-state proliferation (see Discussion).

In addition to metabolic regulators, glial and neuronal genes were required for Q->P. The axon-glial regulator *nrv2* was required for Q->P in Layer 2 FSCs, but not Layer 1 or Layer 0 (Fig. 6, Table S1). We previously demonstrated that Layer 2 FSCs extend long cytoplasmic projections that contribute to the cell fate decision to remain as an FSC or differentiate^37^. As *nrv2* functions as a beta subunit of the sodium potassium ATPase that maintains ion gradients to control neuronal and glial responses^64^, the identification of *nrv2* as a critical regulator of Layer 2 FSC proliferation induction is enticing. To pursue this further, we tested a second beta subunit of the sodium potassium ATPase, *nrv1*, which is generally muscle-specific^64^. *nrv1* was expressed at low levels in *10930*-Gal4 expressing cells, with slight, but significant, feeding dependence based on RNAScope (Fig. 5E,F). Complementary to the requirement for *nrv2* in Layer 2 FSCs, *nrv1* was required in Layer 1 FSCs for proliferation (Fig. 6, Table S1), supporting a model in which Layer-specific ATPase function contributes to proliferation regulation. Multiple regulators of the neurotransmitter glutamate were streptavidin-precipitated and feeding-dependent, including *Ekar, Eaat1, Eaat2*, and *repo* (Fig. 2). In addition, the *GABA Transporter, Gat*, which is expected to reduce GABA signaling, thus indirectly enhancing glutamate signaling, was identified as feeding-dependent. Functionally, only *Gat* was necessary for Q->P in Layer 1 FSCs (Fig. 6, Table S1), suggesting a role for GABA inhibition in Q->P regulation. Further analysis of the role of glutamate in regulation of other cellular processes within FSCs may be warranted, given the enrichment for this pathway in our TU-tagging results (Fig. 2). Together, these results suggest that location-based neuronal-glial signaling may differentially control cell cycle entry or progression in Layer 2 versus Layer 1 FSCs.

## Discussion

Quiescence to proliferation transitions are critical for the long-term health of stem cell populations^65-68^. Both states are regulated extensively, with external signals controlling internal cellular events such as metabolism, gene expression, cell-cell communication, autophagy, and cell cycle status. The balance between quiescent and proliferative states varies among different stem cell populations, but disruption of the balance accelerates stem cell aging and is often associated with inflammation and injury^65-68^. Despite the critical importance of Q->P transitions, the mechanisms that control transitions into and out of quiescence are not fully understood. Cell cycle regulators are the best known regulatory mechanism, with the balance of cyclins, cyclin-dependent kinases (CDKs), and CDK inhibitors determining cell cycle status^69, 70^. Levels of these regulators are controlled via transcription, translation, and post-translational modifications, processes that are regulated by external and internal signals that link cell cycle regulation to the immediate needs of the tissue or organism^69, 70^. Previously, we demonstrated that Hh-mediated transcription is essential for Q->P in FSCs^23, 37^, with an expectation that expression of cyclins, CDKs, or CDKis would be controlled by feeding-dependent induction of Hh signaling. Instead, we found no changes in any known cell cycle regulator (Fig. S2), suggesting that other mechanisms are critical for Q->P in this stem cell population. We found 7 new regulators of Q->P using a multi-step, forward screening approach centered on the gene expression changes that occur in FSCs during the 6-hour Q->P transition. Components of cholesterol regulation (Npc2h) are not unexpected, given prior work demonstrating the critical importance of cholesterol in Hh pathway regulation, including for FSCs^23, 71^. Identification of neuronal and glial regulators highlights the potential functional significance of long axon-like cytoplasmic extensions that characterize FSCs^37^, uncovering new models for testing how ion gradients and GABA function as signaling molecules that contribute to FSC activation in response to feeding. Finally, offsetting the balance between the feeding response hormones NPF and Akh prevents Q->P (Fig. 6, Table S1), suggesting that regulation of macronutrient metabolism via transcriptional regulation is a gatekeeper of the quiescent state. These results highlight the power and potential of forward genetic screening for discovery of novel regulatory mechanisms, with FSCs providing a robust system for analysis of novel mechanisms that regulate important stem cell states.

### Temporal/spatial gene expression screening uncovers novel mechanisms

A major challenge for delineating Q->P regulatory mechanisms in other stem cell populations is technical. Isolation of single cells for RNA sequencing depends on cellular disruption. Injury, changes in cell-cell or cell-matrix adhesion, removal from signaling gradients, or other abrupt manipulations alters stem cells almost immediately^72-74^, making it difficult to interpret changes observed as intrinsic to stem cells or a result of the experimentation. TU-tagging presents a solution^40, 41^. Transcripts are tagged under normal developmental conditions without requiring isolation of individual cells, sorting, or other manipulations that alter the gene expression environment^40, 41^. Whole tissue is isolated and lysed prior to mRNA isolation, freezing the transcriptome at the point of lysis and preventing *ex vivo* changes from occurring^40, 41^. The approach enables analysis of gene expression changes over time, in response to stimuli such as feeding, and in genetic conditions that can test directly signaling or mechanistic hypotheses. Most importantly, TU-tagging does not rely on reference markers or modeling for cell identification. Only cells expressing UPRT can generate TU-tagged transcripts^40, 41^, enriching the precipitated pool of mRNA for transcripts expressed in cells of interest. The approach enables reading at more depth than single cell RNA sequencing, with identification of low-expressed transcripts increasing the potential for discovery of novel regulators of high functional value. Our proof-of-principle TU-tagging study, followed by RNA in situ hybridization and functional analysis, demonstrates that sequential screening can successfully identify new Q->P regulators. In addition to the 46 known genes screened here, 97 known genes and 156 uncharacterized genes were also identified as feeding-induced, candidate FSC regulators, providing an ample pool of genes for further investigation. In addition, genes identified via this approach may be important for FSC functions independently of Q->P, providing a new pool of candidate genes for analysis.

### Nervous tissue markers function in FSCs

Our TU-tagging screen uncovered a striking number of nervous tissue markers as FSC-enriched and feeding dependent. The largest group of these are glial markers, including genes that are used to define all glia (*repo*) or subsets of glia (*moody, CG5080, wrapper, Eaat2, nrv2, alrm, Gat, Eaat1, DAT*). We investigated several of these genes further via RNAScope and functional analysis, finding multiple genes that visibly increased in FSCs after feeding (Fig. 5). *nrv2* and *Gat* were also necessary for feeding-induced Q->P (Fig. 5, 6, Table S1). Glia are critical for homeostasis of the nervous system, functioning to insulate neuronal axons, accelerate signaling, control metabolism, transfer nutrients to neurons, provide immune function, maintain the blood-brain barrier, and dynamically regulate synapses^75^. In mammals, radial glia serve as progenitors that can produce neural stem cells or glia, depending on the developmental stage and environmental context^76^. In *Drosophila*, a subtype of glial cells known as “cortex glia” function as a niche for neural stem cells, both controlling neural stem cell division and protecting and positioning newly generated neurons^77^. Notably, dietary nutrients stimulate Q->P transitions in neural stem cells via glia^78^. Specifically, feeding stimulates glial membrane growth, enabling the physical contact between glia and quiescent neural stem cells that triggers their activation^78^. An exciting possibility is that FSCs use similar mechanisms to wake up stem cells in response to feeding. Our previous work demonstrated dramatic extension of cytoplasmic projections upon feeding, with rapid growth of projections enabling interaction between FSCs located many cell diameters apart to form contacts^79^. Based on analogy with glia-neural stem cell interactions, it is possible that feeding-dependent contact between FSCs stimulates Q->P. Interestingly, prior studies have demonstrated that increased intracellular potassium can trigger G0 or G1 to S transitions, a process that depends on action of sodium-potassium ATPases^80, 81^. Transcriptional induction of *nrv2*, a surface-expressed sodium-potassium ATPase, may enable influx of potassium upon FSC-FSC contact, promoting exit from quiescence. Manipulation of the neurotransmitter GABA may also contribute. In mammals, GABA promotes quiescence of neural stem cells; ablation of GABAergic signaling is required for Q->P ^82^. We found that the GABA transporter, *Gat*, was required for Q->P in Layer 1 FSCs (Fig. 6, Table S1), consistent with its function in clearing GABA from synapses in neurons and glia. Together, the reduction in quiescence-promoting GABA signaling and increased intracellular potassium that occur upon transcriptional induction of *nrv2* and *Gat* may help trigger Q->P in FSCs.

In addition to glial markers, neuronal regulators were identified as important for Q->P in FSCs (Fig. 6, Table S1). We identified *Akhr*, a G-protein coupled receptor for adipokinetic hormone (Akh), as a feeding-induced gene that is required for Q->P. Akh, produced by neuroendocrine cells in a region of the fly brain called the corpora cardiaca, activates Akhr expressed on the surface of cells in the fat body to facilitate breakdown of carbohydrates and lipids in low energy environments, such as in starved conditions^83, 84^. As a functional analog of mammalian glucagon^83, 85, 86^, Akhr function is critical during nutrient restriction, and loss-of-function mutation of *Akhr* results in obesity due to failure to breakdown storage nutrients^87^. In addition to fat body expression, Akhr is expressed on gustatory neurons in the brain, where Akh signaling modulates the fly’s response to hunger by increasing its appetite^85^, thus promoting replacement of spent storage nutrients. Our TU-tagging results demonstrate strong enrichment of *Akhr* in *109-30* Gal4-expressing cells relative to Input RNA, with highest levels observed in nutrient-restricted flies (Fig. 2E). This would be consistent with a function analogous to that observed in the fat body, with Akhr promoting catalysis of lipids and carbohydrates to maintain FSC survival during quiescence. The requirement for *Akhr* for Q->P is less expected. In the fat body and gustatory neurons, feeding suppresses Akhr signaling and consequent nutrient catalysis. By contrast, we found that *Akhr* is necessary for feeding-dependent Q->P in FSCs, suggesting that lipid and/or carbohydrate catalysis is necessary for the transition. Perhaps Akhr leverages mechanisms used in gustatory neurons to promote FSC “appetite”, helping to ensure that sufficient nutrients are secured by FSCs as they transition to a proliferative state.

Complementary to *Akhr*, we found that expression of *Neuropeptide F* (*NPF*) and its receptor, *NPFR* increase after feeding, and both are required for Q->P in FSCs (Fig. 2, 5, 6, Table S1). Whereas the predominant function of Akhr is in nutrient mobilization under nutrient-restricted conditions, NPF is released by the gut in high glucose or lipid environments^88^. The NPF ligand then stimulates NPFR in the brain, suppressing expression of Akh ligand in corpus cordiaca cells and stimulating production of insulin-like peptides to promote lipid anabolism and storage^89^. Loss of NPF leads to catastrophic changes in sugar and lipid metabolism^89^, highlighting its essential role in the feeding response. Thus, NPF opposes Akh function in a metabolic sense. In terms of appetite control, however, NPF functions similarly to Akhr, with high levels of NPF promoting food searching and feeding^90-92^. For FSCs, the functions of NPF and Akhr may modulate the metabolic environment, establishing an optimal balance between catabolism and anabolism that permits Q->P. As such, *Akhr* and *NPF* expressed autonomously by FSCs may control transition states between starving and feeding, with limited catalysis of lipids, triglycerides, or glycogen facilitating the metabolic, gene expression, and behavioral changes necessary for feeding-dependent transitions. Interestingly, NPF is also produced by the gut in response to mating, driving proliferation of germline stem cells in the ovary via NPFR^93^, supporting the notion that NPF triggers proliferation of reproductive stem cells. Potential mechanisms that link NPF signaling directly to cell cycle regulation or indirectly via metabolic regulation as well as collaboration between NPF and Akh signaling are important open questions that may shed light on our understanding of stem cell quiescence to proliferation transitions in general.

### Promoting quiescence

We anticipated the possibility that loss of quiescence-promoting genes might lead to aberrant FSC proliferation under nutrient-restriction conditions, similar to that observed upon knockdown of *ptc* (Fig. 1, Table S1)^23^. Surprisingly, only knockdown of *nrv2*, a subunit of the sodium-potassium ATPase, resulted in significantly increased proliferation of Layer 1 FSCs under nutrient-restricted conditions (p<0.05, Fig. 6, Table S1). *nrv2* also was required for Q->P in Layer 2 FSCs after feeding, and its family member *nrv1* was required for Q->P in Layer 1 FSCs (Fig. 6). Clearly, regulation of intracellular potassium is a critical element of Q->P, and perhaps P->Q transitions, that will require further investigation.

### Cholesterol regulation and Q->P

Cholesterol is the critical trigger of Q->P in FSCs, acting indirectly via activation of the steroid hormone receptor, DHR96 to trigger Hh release and stimulation of Hh signaling with FSCs^23^. In addition, cholesterol is an important regulator of Smo, a key downstream effector of Hh signaling, which is directly modified by cholesterol^71^. Mechanistically, cholesterol activates Smoothened (Smo) directly by binding its extracellular cysteine-rich domain (CRD), a key step for transducing the Hh signal across the membrane^94^. In the absence of Hh ligand, Ptc suppresses Smo by limiting availability of accessible membrane sterols, including cholesterol, keeping the pathway off^94^. When Hh ligand binds to Ptc, its inhibition is relieved, allowing cholesterol and other sterols to accumulate locally and bind to Smo, activating downstream signaling^94^.

Given the importance of cholesterol in Hh signaling and the requirement for Hh in Q->P induction, it stands to reason that cholesterol regulators might be activated to facilitate this process in FSCs. Here, we identified Npc2h as a new downstream effector of Hh that is necessary for Q->P (Fig. 6, Table S1). Niemann–Pick C (NPC1 and NPC2) proteins mediate the transport of cholesterol from endosomes and lysosomes to the ER and other cellular membranes^95^, providing the cholesterol necessary for proper Hh signaling and other cellular functions^96^. Mutations in NPC1 or NPC2 cause Niemann–Pick type C disease, an autosomal recessive disorder characterized by impaired cholesterol trafficking and consequent abnormalities in tissue morphogenesis that impact brain development and motor function, in particular^95, 97^. For FSCs, Npc2h may modulate Hh signaling amplitude directly, via cholesterol modification of Smo, generation of sterols that influence Ptc function, or by impacting cholesterol moieties on Hh ligand. Other cholesterol-dependent mechanisms also may contribute. In the simplest model, cholesterol is necessary for generation of new lipid bilayers, a critical need during cell division. Npc2h may deliver cholesterol to areas of bilayer formation, supporting FSC division. More mechanistically, cholesterol stimulates proliferation in multiple stem cell populations, driving G1-S transitions^98^. Cholesterol levels impact signaling pathways, including Receptor Tyrosine Kinases, Notch, Wnt, Bone Morphogenetic Protein (BMP) and TOR signaling, in addition to Hh, with integration of the various pathways controlling the balance between proliferation and differentiation in stem cell pools^98^. The same panel of signals controls FSC proliferation^9, 14, 23, 29, 36, 39, 47, 99, 100^, highlighting the likely influence of cholesterol on key FSC regulatory signals. Incredibly, cholesterol release is the key trigger for exit from the quiescent dauer phase in *C. elegans*^101^. Degradation of sterol binding proteins releases cholesterol, which is converted into the ligand for a steroid hormone receptor that drives the developmental switch^101^. Although no steroid hormone receptors were identified as feeding dependent in our TU-tagging screen, *Drosophila Hormone Receptor 3* (*DHR3*) was the most enriched gene in our streptavidin-precipitated fraction. This raises the possibility that cholesterol transport via Npc2h enables activation of DHR3 by a steroid derivative, activating this orphan receptor to promote proliferation in a manner similar to that used during the dauer to growth transition in *C. elegans*^101^. The implication of NPC2 itself in mammalian stem cell proliferation control^102, 103^ highlights the likely broad conservation of its role in FSC regulation, with this genetically tractable system providing opportunities for precise delineation of its mechanism of action.

## Acknowledgments

We thank Chris Doe (HHMI, University of Oregon) and Mike Clearney (University of California at Merced) for invaluable guidance on the TU-tagging protocol, and Josh Mell (Drexel University) for RNA sequencing. This work would not have been possible without FlyBase, supported by NIH [5U24HG013300-02]. We also thank *Drosophila* resource centers at Bloomington [NIH P40OD018537], Vienna (VDRC, www.vdrc.at), and Kyoto (DGRC, Kyoto Institute of Technology), and the Developmental Studies Hybridoma bank (NICHD and University of Iowa). This work was supported by NIH grants to E. Lee and D. Zinshteyn (T32 CA009035), A. O’Reilly (R01 HD065800, R21 HD105295), FCCC (P30 CA06927), and by the Kicking Cancer Foundation.

## Methods

### Fly preparation

All fly stocks were raised on standard fly food (7.5 g/l agar, 83.6 g/l cornmeal, 50 ml/l molasses, 20 g/l yeast, 5.2 ml/l propionic acid, 10 ml tegosept/l). Nutrient restriction was accomplished by placing flies in collection cages on grape juice plates (50% grape juice, 1% acetic acid, 3% Bacto-Agar, 0.1% methylparaben in water;^104^ for a minimum of 3 days^23^. Note that molasses plates do not induce quiescence in FSCs^23, 62, 105^ and are thus not appropriate for nutrient restriction conditions needed to analyze Q→P transitions. Re-feeding during the 24-h timecourse was done by adding a water-based paste of baker’s yeast in water to grape juice plates. Flies were maintained at 22°C during crosses and then shifted to 29°C, the permissive temperature for inactivating Gal80ts, to enable Gal4-dependent induction of UAS-controlled gene expression.

### Fly strains and genetics

The following stocks were obtained from the Bloomington Drosophila Stock Center (BDSC, Bloomington, IN), *109-30-Gal4* ^47^ [*y*^*1*^*w*;P(GawB)109-30/CyO*], *ci RNAi* (BDSC31320: *y*^*1*^*v*^*1*^; *P{TRiP.JF01271}attP2*, BDSC31321: *y*^*1*^*v*^*1*^; *P{TRiP.JF01272}attP2*, BDSC31236: *y*^*1*^*v*^*1*^; *P{TRiP.JF01744}attP2), ptc RNAi* (BDSC55686: *y*^*1*^*sc*v*^*1*^*sev*^*21*^; *P{TRiP.HMC03872}attP40), w*^*1118*^ (BDSC3605), *arm-Gal4* (BDSC1560: *w; P{Gal4-arm.S}11*), *pCOG-Gal4* (*otu-Gal4*^*60*^), *cas-Gal4* (BDSC39575: *w*^*1118*^; *P{GMR71C09-Gal4}attP2*)), *Wnt4-Gal4* (BDSC67449: *y*^*1*^*w; Mi{Trojan-Gal4.2}Wnt4*^*MI03717-TG4.2*^), *wrapper-Gal4* (BDSC45784: *w*^*1118*^; *P{y*^*+*^*w*^*+*^=*GMR54H02-Gal4}attP2*), *CG6718-Gal4* (BDSC17388: *y*^*1*^*w*^*67c23*^; P{ *y*^*+*^*w*^*+*^*=EPgy2}CG6178*^*EY07693*^), *CCHa2-Gal4* (BDSC84602: *w*; TI{2A-Gal4}CCHa2*^*2A-Gal42*^), *hkb-Gal4* (BDSC62578: *w*^*1118*^; *PBac{w*^*+*^*=IT.Gal4}hkbGal4*^*0016-G4*^), *toy-Gal4* (BDSC41336: *w*^*1118*^; *P{y*^*+*^*w*^*+*^*=GMR9G09-Gal4}attP2*), *Ubx-Gal4* (BDSC45575: *w*^*1118*^; *P{y*^*+*^*w*^*+*^*=GMR31E11-Gal4}attP2*), *Akhr-Gal4* (BDSC46156: *w*^*1118*^; *P{y*^*+*^*w*^*+*^*=GMR18A10-Gal4}attP2*), *DopEcR-Gal4* (BDSC48047: *w*^*1118*^; *P{y*^*+*^*w*^*+*^=*GMR23D10-Gal4}attP2*), *nrv2-Gal4* (BDSC6794: *w*^*\**^; *P{*^*+*^*w*^*+*^*=nrv2-Gal4.S}8 P{*^*+*^*w*^*+*^*=UAS-GFP.S65T}eg*^*T10*^), *DAT-Gal4* (BDSC39108: *w*^*1118*^; *P{y*^*+*^*w*^*+*^*=GMR55C10-Gal4}attP2*), *alrm-Gal4* (BDSC67031: *w*^*\**^; *P{ w*^*+*^=*alrm-Gal4.D}3*), *Arr1-Gal4* (BDSC83266: *y*^*1*^*w*; TI{GFP[3xP3.cLa]=CRIMIC.TG4.1}Arr1*^*CR01322-TG4.1*^), *Prat-Gal4* (BDSC90994: *w*^*\**^; *P{w*^*+*^*=Prat2-Gal4.10.1}6*), *Lsp-Gal4* (BDSC98128: *w*^*\**^; *P{w*^*+*^*=3.1Lsp2-Gal4}3*), *Ekar-Gal4* (BDSC93679: *y*^*1*^*w*^*1118*^; *Mi{DH.1}EkarMI02500-DH.GT-TG4.1}*), *Npc2h-Gal4* (BDSC98630: *y*^*1*^*w*^*1118*^; *TI{GFP*^*3xP3*.cLa^=*KozakGal4}Npc2h*^*CR70738-KO-kG4*^*}*), *NPF-Gal4* (BDSC25681 *y*^*1*^*w*^*\*8*^; *P{w[+mC]*=*NPF-Gal4.1}2*), *ap-Gal4* (BDSC3041: *y*^*1*^*w*^*1118*^; *P{w*^*+*^*=GawB}ap*^*md544*^), *Odc1-Gal4* (BDSC66812: *y*^*1*^*w; Mi{Trojan-Gal4.0}Odc1*^*MI10996-TG4.2*^), *Ndg-Gal4* (BDSC76768: *y*^*1*^*w; Mi{Trojan-Gal4.0}Odc1*^*MI15397-TG4.2*^), *NimB5-Gal4* (BDSC86378: *y*^*1*^*w*; TI{GFP[3xP3.cLa]=CRIMIC.TG4.2}NimB5^CR01426-TG4.2^*), *ninaB-Gal4* (BDSC24518: *w*^*\**^; *P{w*^*+*^*=ninaB-Gal4.W}2*), *CrebA-Gal4* (BDSC46512: *w*^*1118*^; *P{y*^*+*^*w*^*+*^*=GMR64A03-Gal4}attP2*), *dsx-Gal4* (BDSC41250: *w*^*1118*^; *P{y*^*+*^*w*^*+*^*=GMR42D02-Gal4}attP2*), *dtr-Gal4* (BDSC62593: *w*^*1118*^; *PBac{w*^*+*^*=IT.Gal4}dtr*^*0043-G4*^), *Rh6-Gal4* (BDSC7464: *w*^*\**^; *P{w*^*+*^*=Rh6-Gal4.D}3*), *Pdh-Gal4* (BDSC19425: *w*^*\**^; *PBac{GAL4D*,*EYPF}Pdh*^*PL00234*^*Tcs3* ^*PL00234*^), *dpr1-Gal4* (BDSC25083: *w*^*\**^; *P{w*^*+*^.*hs=GawB}dpr1*^*PGaw*^), *Lkr-Gal4* (BDSC39261: *w*^*1118*^; *P{y*^*+*^*w*^*+*^*=GMR60G10-Gal4}attP2*), *dtr-Gal4* (BDSC86463: *y*^*1*^*w*; TI{GFP[3xP3.cLa]=CRIMIC.TG4.2}dtr*^*CR01606-TG4.2*^), *Clk-Gal4* (BDSC41259: *w*^*1118*^; *P{y*^*+*^*w*^*+*^*=GMR43D05-Gal4}attP2*), *Actbeta-Gal4* (BDSC49035: *w*^*1118*^; *P{y*^*+*^*w*^*+*^*=GMR23F05-Gal4}attP2*), *Gr59f-Gal4* (BDSC57656: *w*^*\**^; *P{w*^*+*^*=Gr59f-Gal4.1.8}6*), *Syp-Gal4* (BDSC66841: *y*^*1*^*w; Mi{Trojan-Gal4.1}Syp*^*MI06413-TG4.1*^), *CCAP-R-Gal4* (BDSC84600: *TI{2A-Gal4}CCAP-R*^*2A-Gal4*^), *Mtp-Gal4* (BDSC86445: *y*^*1*^*w*; TI{GFP[3xP3.cLa]=CRIMIC.TG4.1}Mtp*^*CR01555-TG4.1*^), *LKRSDH-Gal4* (BDSC92724: *y*^*1*^*w*; TI{GFP[3xP3.cLa]=CRIMIC.TG4.0}LKRSDH*^*CR70173-TG4.0*^), *Tpst-Gal4* (BDSC97575: *y*^*1*^*w*; TI{GFP[3xP3.cLa]=CRIMIC.TG4.1}CG4404*^*CR02713-TG4.1*^), *GABA-B-R1-Gal4* (BDSC84701: *w*; TI{2A-Gal4}GABA-B-R1*^*2A-Gal4*^), *apolpp-Gal4* (BDSC97179: *y*^*1*^*w*; TI{GFP[3xP3.cLa]=CRIMIC.TG4.1}apolpp*^*CR70471-TG4.1*^), *moody-Gal4* (BDSC90883: *w*^*\**^; *P{w[+m*]*=*moody-Gal4.SPG}2*), *nrv1-Gal4* (BDSC76203: *y*^*1*^*w; Mi{Trojan-Gal4.1}nrv1*^*MI094257-TG4.1*^), *Akhr-RNAi (BDSC29547*: *yv;P{y*^*+*^*v*^*+*^*=TRiP.JF02439}attP2*), *ap-RNAi (BDSC26748: yv;P{y*^*+*^*v*^*+*^*=TRiP.JF02311}attP2), Arg-RNAi (BDSC56881: y*^*1*^ *sc* v*^*1*^ *sev*^*21*^*;P{y*^*+*^*v*^*+*^*=TRiP.HMC04102}attP40/CyO), Ekar-RNAi (BDSC41832: yv;P{y*^*+*^*v*^*+*^*=TRiP.GL01260}attP2/TM3), Gat-RNAi (BDSC29422: yv;P{y*^*+*^*v*^*+*^*=TRiP.JF03358}attP2), Npc2h-RNAi (BDSC67808: y*^*1*^ *sc* v*^*1*^ *sev*^*21*^*;P{y*^*+*^*v*^*+*^*=TRiP.HMS05621}attP40), NPF-RNAi (BDSC27237: yv;P{y*^*+*^*v*^*+*^*=TRiP.JF02555}attP2), NPFR-RNAi (BDSC25939: yv;P{y*^*+*^*v*^*+*^*=TRiP.JF01959}attP2), nrv1-RNAi (BDSC38275: y*^*1*^ *sc* v*^*1*^ *sev*^*21*^*;P{y*^*+*^*v*^*+*^*=TRiP.HMS01725}attP40), nrv2-RNAi (BDSC28666: yv;P{y*^*+*^*v*^*+*^*=TRiP.JF03081}attP2)*.

### Dissections and immunostaining

For all experiments, 15-60 female flies of the indicated genotype were selected at random from a larger pool. Ovaries from ~1-week-old adult female flies (*Drosophila melanogaster*) were dissected in Grace’s insect cell culture medium (Gibco, Gaithersburg, MD, USA), fixed in 4% paraformaldehyde for 15 min and then washed three times in 1X PBST for 5 min. The ovaries were then incubated with primary antibodies in 0.5% BSA diluted with 1X PBST solution overnight at 4°C. The ovaries were washed three times for 10 min each in 1X PBST and then incubated with secondary antibodies overnight at 4°C. Ovaries were washed three times in 1X PBST. For nuclei labeling, ovaries were incubated in eBioscience Draq5 (Thermo-Fisher, Waltham, MA^106^) for 10 min at room temperature. The ovaries were then mounted on slides using Vectashield medium (Vector Laboratories, Burlingame, CA, USA).

### Reagents

Primary antibodies used were mouse anti-Fasciclin III (FasIII) (1:200; 7G10, DSHB, Iowa City, IA^107^), rabbit anti-PH3 (1:1000, Cat# 06-570, MilliporeSigma). Secondary antibodies used were Rhodamine-conjugated Goat anti-Mouse (Cat# 115-025-003) and fluorescein-conjugated Goat anti-Rabbit (Cat# 111-095-144) (1:200;Jackson ImmunoResearch Laboratories). Draq5 was used to label nuclei (1:1000, Miltenyi Biotec Cat# 130-117-344).

RNAScope probes against Gal4 were available from ACD Bio (Cat# 428501). Drosophila Fas3 probes were custom designed for this project and are now available (Cat# 1323891-C4). Detection reagents for C1-linked (Gal4, Cat # 323110) or C4-linked (Fas3, Cat# 323120) probes were from ACD Bio.

### TU-tagging and RNA sequencing

25-30 bottles of flies were cultured, and females were collected at 7 days after eclosion. Flies were starved on grape juice bottles for 2 days, and subsequently transferred into an empty bottle (no food no water) for 12 hours before feeding flies with 1mM 4TU (Sigma 440736) in 10% sucrose solution soaked in cotton. Flies were fed with 4TU sucrose solution alone for 24 hours and then fed with yeast paste containing 1mM 4TU. Flies were collected in Graces medium and blended with a high pulse blender at the lowest setting. Solutions containing ovaries were filtered through a 250um nylon mesh. The collected ovaries were then suspended in 5-7mL Trizol and sheared with a 25G syringe. RNA was isolated, and TU tagged RNA from starved, 3, 6, or 24 hr fed flies was biotinylated and immunoprecipitated by streptavidin beads. Concentration was measured by Ribogreen RNA assay (Thermo R11490).

Data was collected from 3 separate RNA pools (biological triplicates) derived from >100 ovary pairs each. Library production was performed by the Genomics Core Facility at Drexel University College of Medicine using the Illumina TruSeq stranded mRNA kit according to recommended protocols. TU-tagged libraries and Input RNA were sequenced to a depth of at least 15M unpaired reads, which were assessed for quality and trimmed as necessary using Cutadapt^108^. Reads were then aligned to the Drosophila melanogaster genome (dm6) using the STAR aligner^109^. HT-seq^110^ was used to assigns aligned reads to genomic features. Raw read counts were imported into DESeq2 ^111, 112^, normalized and tested for differential expression via negative binomial generalized linear models that incorporated time and/or genotype. Benjamini-Hochberg method^113^ was applied to adjust for multiple comparisons, controlling the false discovery rate (FDR). Genes with a fold change of at least two relative to paired wild-type samples and a false discovery rate of less than 5% (FDR < 0.05) were considered statistically significant.

### Data Availability

Raw and processed RNA-sequencing data have been deposited in the NCBI Gene Expression Omnibus (GEO) under accession number **GSE317804**.

### RNAScope

Female flies from starve-feed and steady-fed experiments were dissected in Grace’s insect cell culture medium and transferred to an Eppendorf tube. Ovaries were fixed in 4% paraformaldehyde and incubated overnight at 4°C. This was followed by a methanol dehydration series (25%, 50%, 75%, 100%) in Phosphate Buffered Saline + 0.3% Triton (PBST) solution, Ovaries were stored in 100% methanol at −20°C until probing. The process for binding of the probe began by transferring the ovaries to a glass dish, removing methanol from the samples, and allowing them to dry. Hydrogen peroxide was then added to each well and removed after an incubation period of 5 minutes. RNAScope Multiplex kit (Advanced Cell Diagnostics) was used for the following steps: Samples were treated with Protease 4 for 5 minutes followed by 3X washing with PBST. Samples were then incubated in a solution composed of Gal4 probe and Fas3 probe at 40°C overnight. After washing 3X in 1X RNAScope wash buffer, ovaries were then fixed in 4% paraformaldehyde at room temperature for 10 minutes and washed 3X in 1x RNAScope buffer. The amplification process entailed adding RNAScope AMP1, AMP2, and AMP3 to each sample and incubating for a period of 30 minutes sequentially, with 3X washes between incubations. Probes were labeled with fluorescent markers by 15–30-minute incubation at 40°C with 1) HRP-C4 followed by TSA vivid 570 to label Fas3 or 2) HRP-C1 followed by TSA Vivid 520 to label Gal4. HRP-Blocker was used between incubations to stop each reaction. After the final wash, Draq5 (Miltenyi Biotec) was added to each well and incubated at room temperature for 10 minutes to label nuclei. Ovaries were then transferred to an Eppendorf tube where the Draq5 was replaced with Vectashield mounting medium (Vector Laboratories) used to preserve the fluorescence when mounted on slides.

### Proliferation assay

Flies were generated by crossing *109-30-Gal4TubGal80*^*ts*^/CyO to their corresponding UAS-RNAi. Flies carrying *109-30-Gal4TubGal80*^*ts*^/UAS-RNAi were incubated at 29°C for three days prior to dissection. All samples were starved for 3 days prior to re-feeding with yeast for corresponding time points. Ovaries were dissected in Grace’s insect medium and stained with rabbit anti-phospho-histone-H3 (PH3), mouse anti-FasIII and Draq5 to label nuclei. After completing the immunofluorescence procedure described above, mitotic index was calculated as the number of germaria with at least one PH3-positive cell in specific Layers (as indicated in the figures), divided by the total number of germaria^47, 114^. Statistical differences were determined using the Student’s *t* test for two samples, with significance achieved at p ≤ 0.05.

### Image analysis and quantitation

Images were collected at room temperature using a 40X (1.25 NA) oil immersion lens (Leica) on an upright microscope (DM 5000; Leica Microsystems, Wetzlar, Germany) coupled to a confocal laser scanner (TCS SP5; Leica). LAS AF SP5 software (Leica) was used for data acquisition. Images representing individual channels of single confocal slices or three-dimensional reconstructions of the germarium, including the FSC region were exported into IMARIS or Fiji (ImageJ) for further analysis. Photoshop (Adobe) was used to compile figures and add ovals highlighting the FSC niche. For semi-quantitative analysis of RNAScope images, regions of interest were drawn around the FSC niche using Fiji. Green fluorescence (Gal4 probe) was calculated for each pixel, then averaged. Averages for each image were compiled with average fluorescence intensity from all images from the same genotype, resulting in a composite average and variance. Significance was calculated using Student t-tests comparing nutrient-restricted versus 6 hour fed conditions from the same genotypes. A minimum of 5 images was used for analysis of each genotype.

## Supplementary Figures

**Figure S1:**
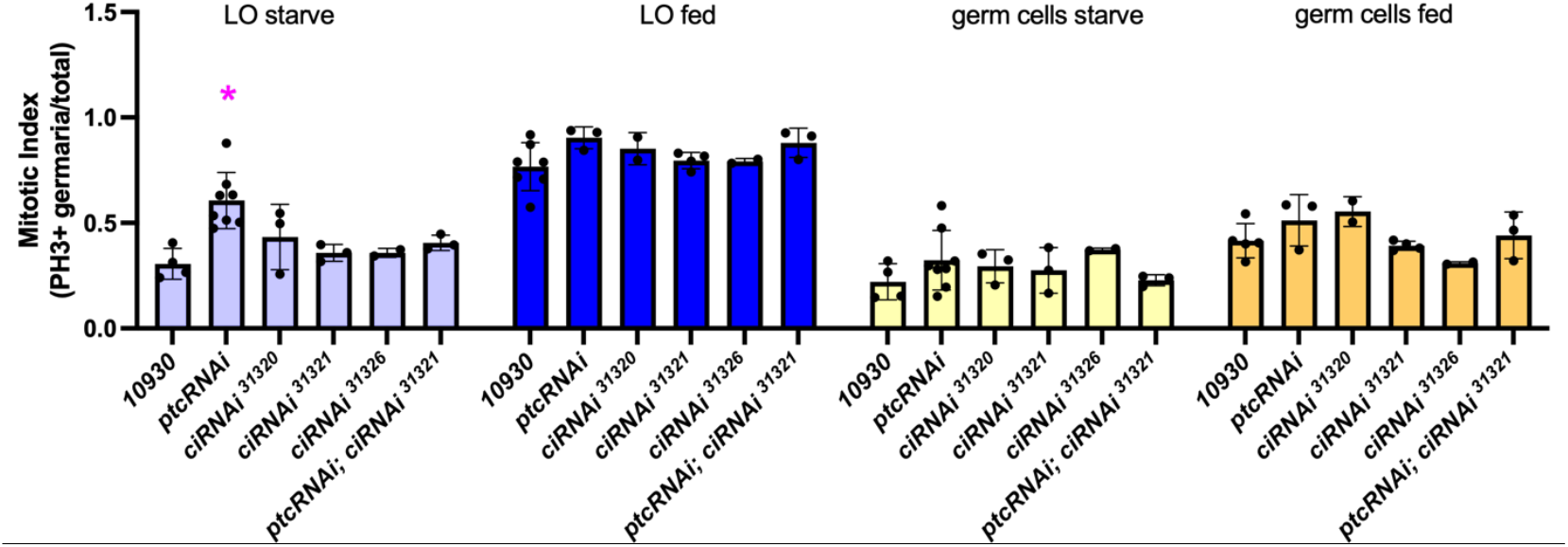
Mitotic index of L0 differentiating follicle cells (blue) or germ cells (yellow) for the indicated genotypes. Lighter colors indicate starved conditions; darker colors indicate fed. * indicates p<0.01 relative to 10930 starved. No other significant differences were observed.

**Figure S2:**
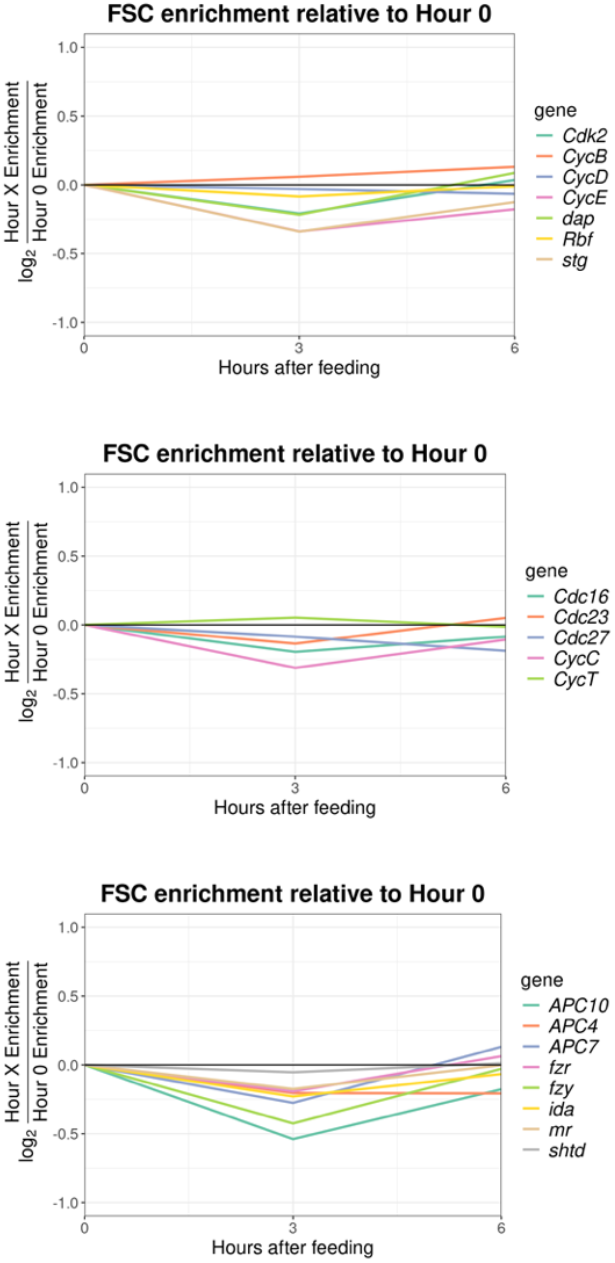
Cell cycle regulatory genes do not change expression upon feeding. Enrichment of TU-tagged cell cycle genes/Input at 3 or 6 hours after feeding relative to enrichment at time 0 (nutrient-restricted). Changes >2-fold and FDR<0.05 were considered significant. No cell cycle regulators changed significantly.

**Figure S3:**
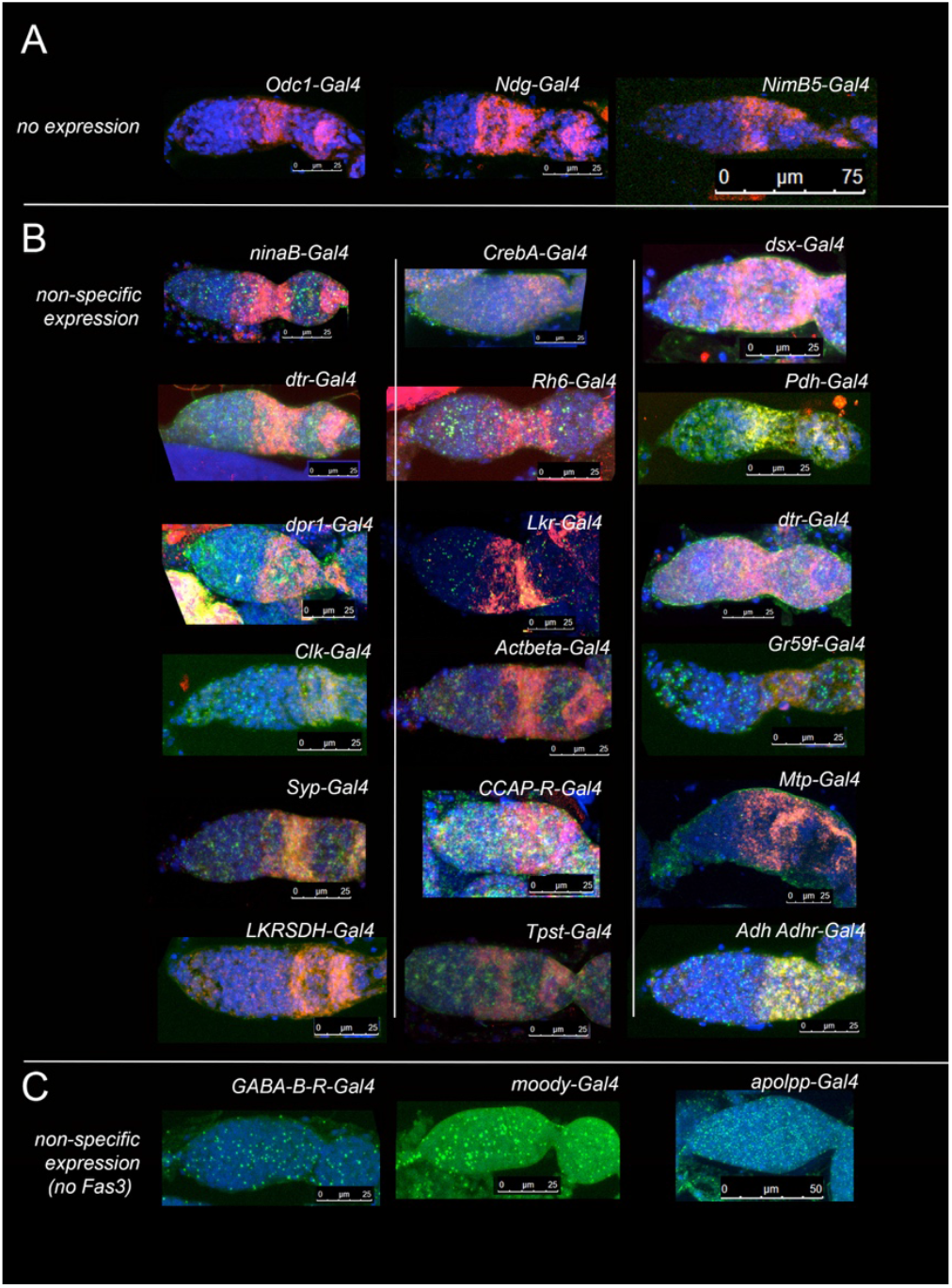
RNAScope of transgenic lines bearing insertion of Gal4 in Q->P regulatory candidates, using probes against Gal4 (green) and Fas3 (red). Nuclei are labeled (Draq5, blue). Candidates with no detectable Gal4 (top) or broad, non-specific expression (middle, bottom) are shown. Scale bars are indicated for each image.

**Table S1:**
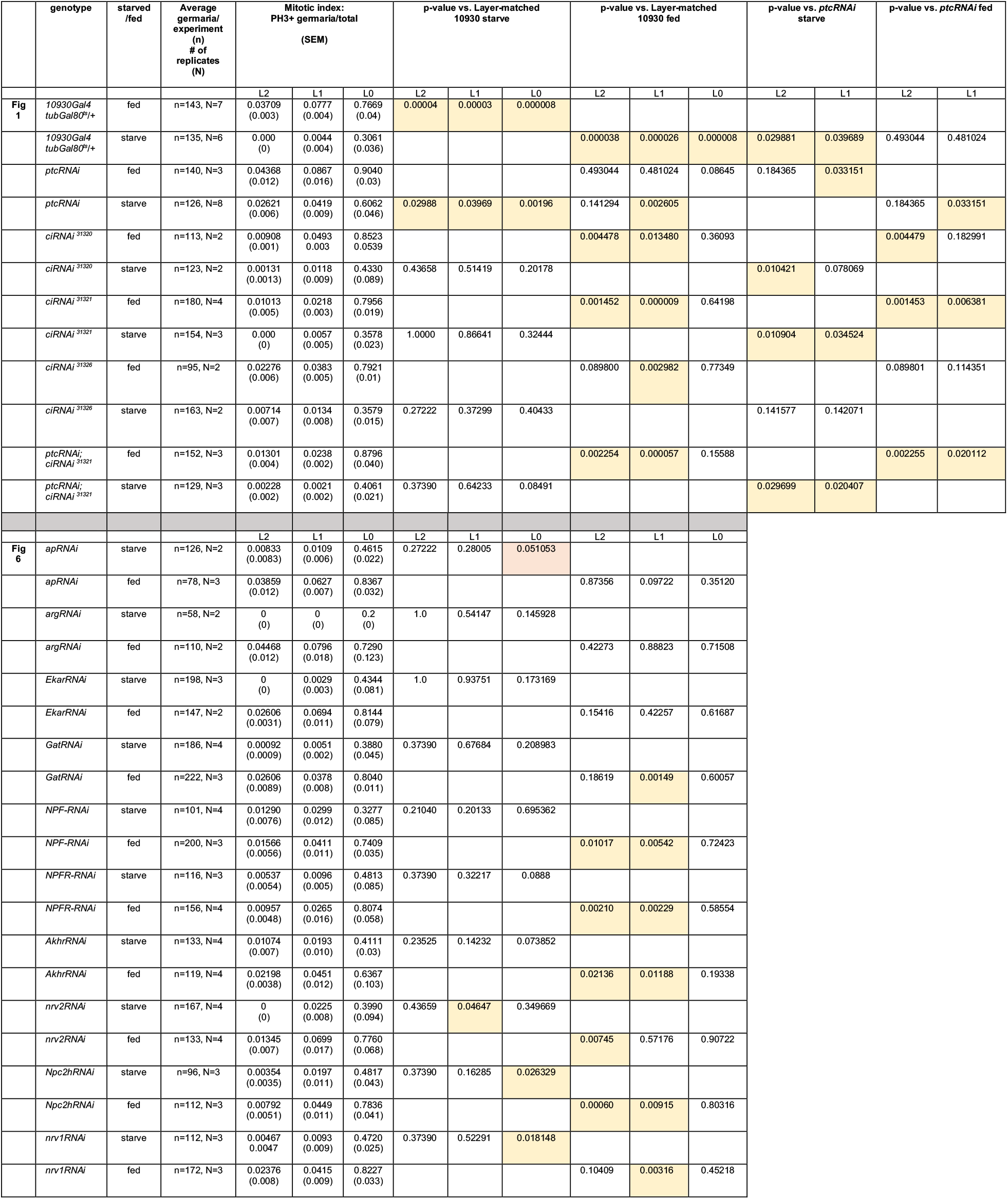
Mitotic index calculations for starve or fed conditions. Unpaired, two-tailed Student t-tests were used for calculation of significance (p<0.05). Significant differences are highlighted in yellow. p=0.05 is highlighted in peach.

## Notes

https://www.ncbi.nlm.nih.gov/geo/query/acc.cgi?acc=GSE317804

### Competing Interest Statement

The authors have declared no competing interest.

### Summary of Updates

Addition of GEO accession number: "Data Availability Raw and processed RNA-sequencing data have been deposited in the NCBI Gene Expression Omnibus (GEO) under accession number GSE317804."

