## Supplementary Figures and Table for "Starve-Feed Cycles Direct Quiescence to Proliferation Transitions in *Drosophila* Follicle Stem Cells via Transcriptional Regulation"

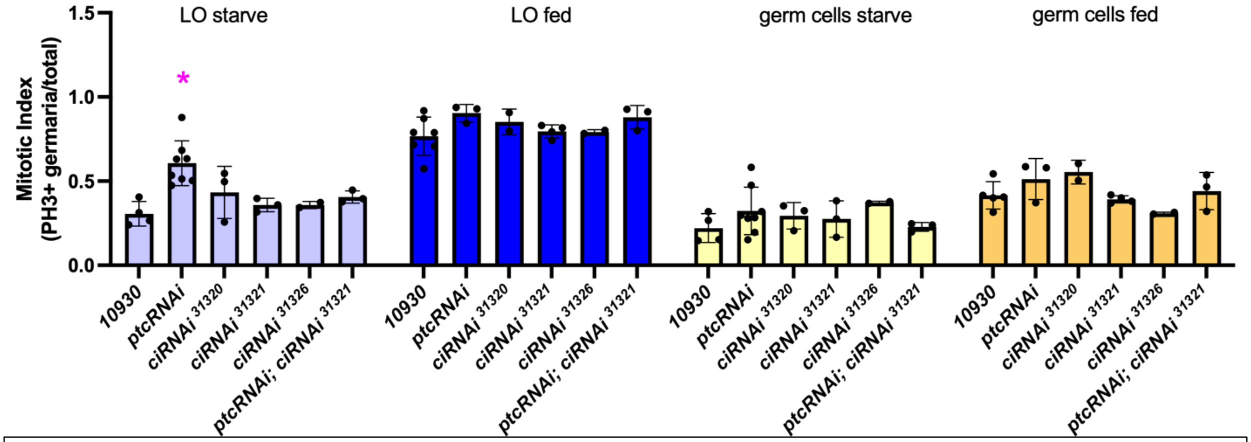

**Figure S1**: Mitotic index of L0 differentiating follicle cells (blue) or germ cells (yellow) for the indicated genotypes. Lighter colors indicate starved conditions; darker colors indicate fed. * indicates $p<0.01$ relative to 10930 starved. No other significant differences were observed.

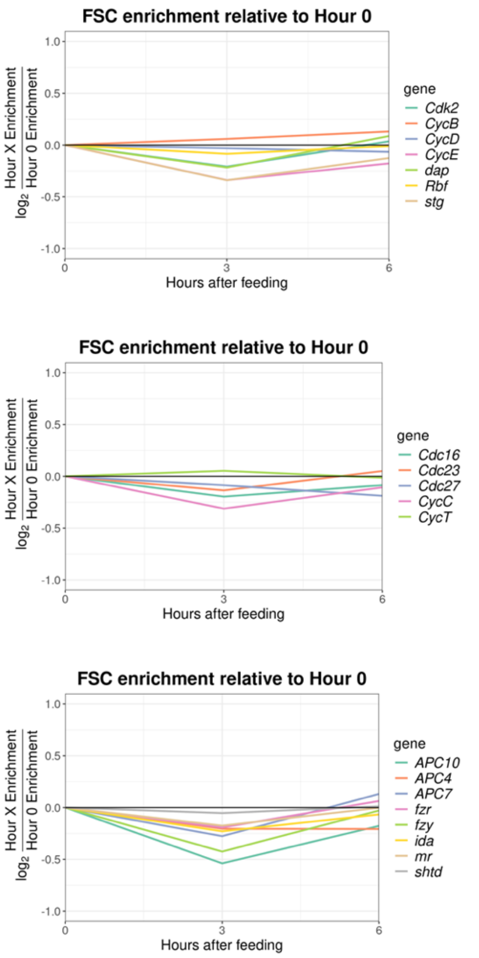

**Figure S2:** Cell cycle regulatory genes do not change expression upon feeding. Enrichment of TU-tagged cell cycle genes/Input at 3 or 6 hours after feeding relative to enrichment at time 0 (nutrient-restricted). Changes >2-fold and FDR<0.05 were considered significant. No cell cycle regulators changed significantly.

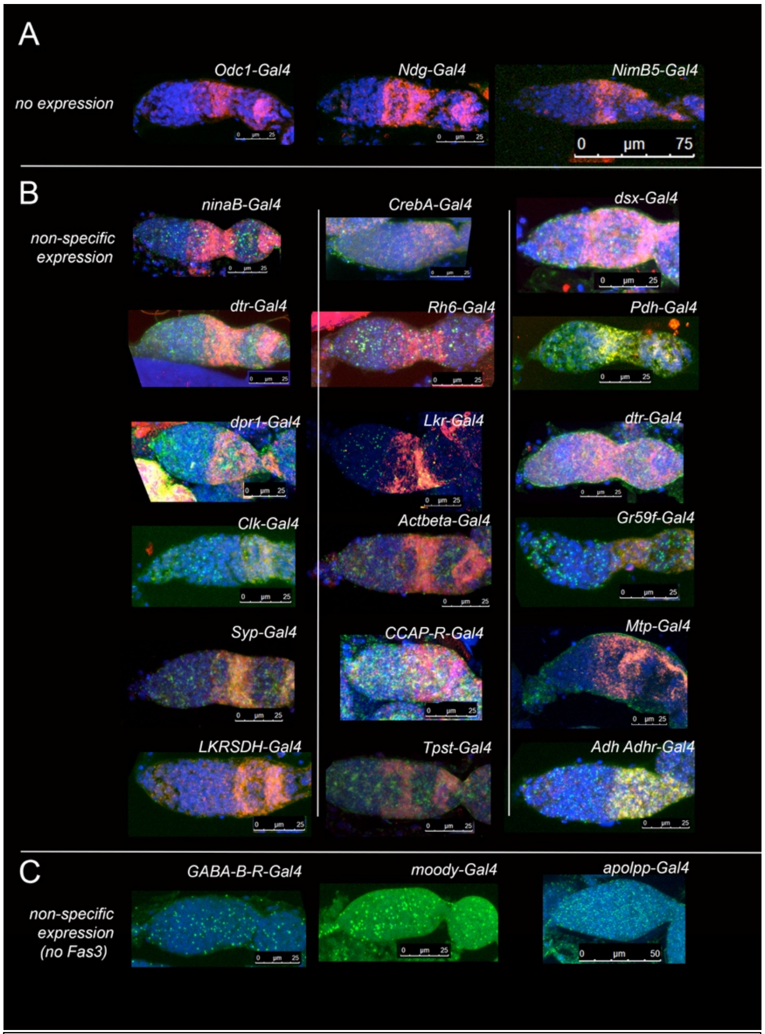

**Figure S3:** RNAScope of transgenic lines bearing insertion of Gal4 in Q->P regulatory candidates, using probes against Gal4 (green) and Fas3 (red). Nuclei are labeled (Draq5, blue). Candidates with no detectable Gal4 (top) or broad, non-specific expression (middle, bottom) are shown. Scale bars are indicated for each image.

| | genotype | starved /fed | Average germaria/ experiment (n) # of replicates (N) | Mitotic index: PH3+ germaria/total (SEM) | | | p-value vs. Layer-matched 10930 starve | | | p-value vs. Layer-matched 10930 fed | | | p-value vs. *ptcRNAi* starve | | p-value vs. *ptcRNAi* fed | |
|---|---|---|---|---|---|---|---|---|---|---|---|---|---|---|---|---|
| | | | | L2 | L1 | L0 | L2 | L1 | L0 | L2 | L1 | L0 | L2 | L1 | L2 | L1 |
| **Fig 1** | *10930Gal4 tubGal80*$^{ts}$/+ | fed | n=143, N=7 | 0.03709 (0.003) | 0.0777 (0.004) | 0.7669 (0.04) | 0.00004 | 0.00003 | 0.000008 | | | | | | | |
| | *10930Gal4 tubGal80*$^{ts}$/+ | starve | n=135, N=6 | 0.000 (0) | 0.0044 (0.004) | 0.3061 (0.036) | | | | 0.000038 | 0.000026 | 0.000008 | 0.029881 | 0.039689 | 0.493044 | 0.481024 |
| | *ptcRNAi* | fed | n=140, N=3 | 0.04368 (0.012) | 0.0867 (0.016) | 0.9040 (0.03) | | | | 0.493044 | 0.481024 | 0.08645 | 0.184365 | 0.033151 | | |
| | *ptcRNAi* | starve | n=126, N=8 | 0.02621 (0.006) | 0.0419 (0.009) | 0.6062 (0.046) | 0.02988 | 0.03969 | 0.00196 | 0.141294 | 0.002605 | | | | 0.184365 | 0.033151 |
| | *ciRNAi* $^{31320}$ | fed | n=113, N=2 | 0.00908 (0.001) | 0.0493 0.003 | 0.8523 0.0539 | | | | 0.004478 | 0.013480 | 0.36093 | | | 0.004479 | 0.182991 |
| | *ciRNAi* $^{31320}$ | starve | n=123, N=2 | 0.00131 (0.0013) | 0.0118 (0.009) | 0.4330 (0.089) | 0.43658 | 0.51419 | 0.20178 | | | | 0.010421 | 0.078069 | | |
| | *ciRNAi* $^{31321}$ | fed | n=180, N=4 | 0.01013 (0.005) | 0.0218 (0.003) | 0.7956 (0.019) | | | | 0.001452 | 0.000009 | 0.64198 | | | 0.001453 | 0.006381 |
| | *ciRNAi* $^{31321}$ | starve | n=154, N=3 | 0.000 (0) | 0.0057 (0.005) | 0.3578 (0.023) | 1.0000 | 0.86641 | 0.32444 | | | | 0.010904 | 0.034524 | | |
| | *ciRNAi* $^{31326}$ | fed | n=95, N=2 | 0.02276 (0.006) | 0.0383 (0.005) | 0.7921 (0.01) | | | | 0.089800 | 0.002982 | 0.77349 | | | 0.089801 | 0.114351 |
| | *ciRNAi* $^{31326}$ | starve | n=163, N=2 | 0.00714 (0.007) | 0.0134 (0.008) | 0.3579 (0.015) | 0.27222 | 0.37299 | 0.40433 | | | | 0.141577 | 0.142071 | | |
| | *ptcRNAi; ciRNAi* $^{31321}$ | fed | n=152, N=3 | 0.01301 (0.004) | 0.0238 (0.002) | 0.8796 (0.040) | | | | 0.002254 | 0.000057 | 0.15588 | | | 0.002255 | 0.020112 |
| | *ptcRNAi; ciRNAi* $^{31321}$ | starve | n=129, N=3 | 0.00228 (0.002) | 0.0021 (0.002) | 0.4061 (0.021) | 0.37390 | 0.64233 | 0.08491 | | | | 0.029699 | 0.020407 | | |
| | | | | L2 | L1 | L0 | L2 | L1 | L0 | L2 | L1 | L0 | | | | |
| **Fig 6** | *apRNAi* | starve | n=126, N=2 | 0.00833 (0.0083) | 0.0109 (0.006) | 0.4615 (0.022) | 0.27222 | 0.28005 | 0.051053 | | | | | | | |
| | *apRNAi* | fed | n=78, N=3 | 0.03859 (0.012) | 0.0627 (0.007) | 0.8367 (0.032) | | | | 0.87356 | 0.09722 | 0.35120 | | | | |
| | *argRNAi* | starve | n=58, N=2 | 0 (0) | 0 (0) | 0.2 (0) | 1.0 | 0.54147 | 0.145928 | | | | | | | |
| | *argRNAi* | fed | n=110, N=2 | 0.04468 (0.012) | 0.0796 (0.018) | 0.7290 (0.123) | | | | 0.42273 | 0.88823 | 0.71508 | | | | |
| | *EkarRNAi* | starve | n=198, N=3 | 0 (0) | 0.0029 (0.003) | 0.4344 (0.081) | 1.0 | 0.93751 | 0.173169 | | | | | | | |
| | *EkarRNAi* | fed | n=147, N=2 | 0.02606 (0.0031) | 0.0694 (0.011) | 0.8144 (0.079) | | | | 0.15416 | 0.42257 | 0.61687 | | | | |
| | *GatRNAi* | starve | n=186, N=4 | 0.00092 (0.0009) | 0.0051 (0.002) | 0.3880 (0.045) | 0.37390 | 0.67684 | 0.208983 | | | | | | | |
| | *GatRNAi* | fed | n=222, N=3 | 0.02606 (0.0089) | 0.0378 (0.008) | 0.8040 (0.011) | | | | 0.18619 | 0.00149 | 0.60057 | | | | |
| | *NPF-RNAi* | starve | n=101, N=4 | 0.01290 (0.0076) | 0.0299 (0.012) | 0.3277 (0.085) | 0.21040 | 0.20133 | 0.695362 | | | | | | | |
| | *NPF-RNAi* | fed | n=200, N=3 | 0.01566 (0.0056) | 0.0411 (0.011) | 0.7409 (0.035) | | | | 0.01017 | 0.00542 | 0.72423 | | | | |
| | *NPFR-RNAi* | starve | n=116, N=3 | 0.00537 (0.0054) | 0.0096 (0.005) | 0.4813 (0.085) | 0.37390 | 0.32217 | 0.0888 | | | | | | | |
| | *NPFR-RNAi* | fed | n=156, N=4 | 0.00957 (0.0048) | 0.0265 (0.016) | 0.8074 (0.058) | | | | 0.00210 | 0.00229 | 0.58554 | | | | |
| | *AkhrRNAi* | starve | n=133, N=4 | 0.01074 (0.007) | 0.0193 (0.010) | 0.4111 (0.03) | 0.23525 | 0.14232 | 0.073852 | | | | | | | |
| | *AkhrRNAi* | fed | n=119, N=4 | 0.02198 (0.0038) | 0.0451 (0.012) | 0.6367 (0.103) | | | | 0.02136 | 0.01188 | 0.19338 | | | | |
| | *nrv2RNAi* | starve | n=167, N=4 | 0 (0) | 0.0225 (0.008) | 0.3990 (0.094) | 0.43659 | 0.04647 | 0.349669 | | | | | | | |
| | *nrv2RNAi* | fed | n=133, N=4 | 0.01345 (0.007) | 0.0699 (0.017) | 0.7760 (0.068) | | | | 0.00745 | 0.57176 | 0.90722 | | | | |
| | *Npc2hRNAi* | starve | n=96, N=3 | 0.00354 (0.0035) | 0.0197 (0.011) | 0.4817 (0.043) | 0.37390 | 0.16285 | 0.026329 | | | | | | | |
| | *Npc2hRNAi* | fed | n=112, N=3 | 0.00792 (0.0051) | 0.0449 (0.011) | 0.7836 (0.041) | | | | 0.00060 | 0.00915 | 0.80316 | | | | |
| | *nrv1RNAi* | starve | n=112, N=3 | 0.00467 0.0047 | 0.0093 (0.009) | 0.4720 (0.025) | 0.37390 | 0.52291 | 0.018148 | | | | | | | |
| | *nrv1RNAi* | fed | n=172, N=3 | 0.02376 (0.008) | 0.0415 (0.009) | 0.8227 (0.033) | | | | 0.10409 | 0.00316 | 0.45218 | | | | |

**Table S1**: Mitotic index calculations for starve or fed conditions. Unpaired, two-tailed Student t-tests were used for calculation of significance ($p<0.05$). Significant differences are highlighted in yellow. $p=0.05$ is highlighted in peach.
